# Mapping Leukocyte Dynamics during Neuroinflammation Identifies Meningeal Monocyte-Derived Macrophages as Drivers of Progressive Disease

**DOI:** 10.1101/2025.08.15.670462

**Authors:** Juan Villar-Vesga, Pauline Clément, Hannah Van Hove, Elèni Meuffels, Viola Bugada, Jeanne Kim, Deborah Greis, Pau Jorba-Adolff, Maud Mayoux, Anna Lasne, Christian Ashworth, Pascale Zwicky, Stefanie Schärli, Hendrik J Engelenburg, Cheng-Chih Hsiao, Jörg Hamann, Joost Smolders, Laura Tatsch, Daniel Kirschenbaum, Bettina Schreiner, Marina Herwerth, Sonia Tugues, Melanie Greter, Donatella DeFeo, Burkhard Becher, Florian Ingelfinger, Sarah Mundt

**Affiliations:** Institute of Experimental Immunology, University of Zurich, Zurich, Switzerland; Department of Neurology, University Hospital Zurich, Zurich, Switzerland; Institute of Pharmacology and Toxicology, University of Zurich, Switzerland; Department of Systems Biology, Weizmann Institute of Science, Rehovot, Israel; Neuroimmunology Research Group, Netherlands Institute for Neuroscience, Amsterdam, The Netherlands; Department of Experimental Immunology, Amsterdam University Medical Centers, Amsterdam, The Netherlands; Departments of Neurology and Immunology, MS Center ErasMS, Erasmus Medical Center, Rotterdam, The Netherlands; Institute of Neuropathology, University Hospital Switzerland, Zurich, Switzerland; Department of Internal Medicine I, Medical Center-University of Freiburg, Freiburg, Germany

**Author notes:** these authors contributed equally to this work (Florian Ingelfinger, Sarah Mundt).

## Abstract

Multiple sclerosis (MS) is a chronic inflammatory disease of the central nervous system (CNS) characterized by increasing disability. The cellular and molecular drivers of clinical transition towards progressive disease are poorly understood. Here, we combine single-cell profiling technologies with genetic and pharmacological perturbations across the course of murine CNS inflammation to dissect the role of the local immune landscape in disease progression. We uncover a chronic monocyte-to-phagocyte transition as a hallmark of progressive disease, characterized by the emergence of maladaptive, lipid-associated macrophages (LAMs) marked by lysosomal activation and fibrotic features. Spatial transcriptomics and multiplexed imaging revealed that these LAMs localized to the leptomeninges in close proximity to parenchymal colony-stimulating factor (CSF)-1 producing disease-associated microglia (DAMs) and meningeal granulocyte-macrophage (GM)-CSF-expressing T helper cells that license their differentiation. Interference with this local cytokine network revealed a protective role for resident microglia and implicated monocyte-derived phagocytes as key drivers of progressive neuroinflammation. Notably, LAM-like macrophages could also be identified in the meninges of people with MS, indicating a homology to human disease. By elucidating their ontogeny, spatial niche, and regulatory cytokine milieu, we provide a mechanistic framework for targeting harmful myeloid states while preserving reparative CNS immunity in progressive MS.

## Introduction

Multiple sclerosis (MS) is a chronic inflammatory disease of the central nervous system (CNS) that often begins with a relapsing-remitting course characterized by episodes of neurological dysfunction followed by full or partial recovery. In many patients, this is eventually followed by a secondary progressive phase (SPMS), marked by a steady accumulation of disability, including worsening of motor and cognitive symptoms with advancing age, with or without continued relapse activity^1^. This progression may begin already during the relapsing phase and may occur independently of relapse activity, reflecting underlying neurodegenerative processes that are not directly linked to focal inflammatory lesions^2^. The gradual transition to SPMS represents a critical shift in disease course, yet remains mechanistically poorly understood, difficult to predict, and challenging to diagnose due to its heterogeneous clinical presentation^3^. Therapeutic options that can halt or reverse neurodegeneration once progression is established are scarce^4^.

Pathological hallmarks of the progressive MS include diffuse microglial activation, meningeal inflammation, slowly expanding lesions, and widespread axonal loss, typically in the absence of overt immune infiltration^5–8^. Increasing evidence implicates CNS phagocytes - resident microglia and infiltrating monocyte-derived cells (MdCs) - as key contributors to this smoldering inflammation and ongoing tissue damage. However, distinguishing these cell types *in situ* has been a central technical challenge, as both resident macrophages as well as MdCs present highly overlapping transcriptional and phenotypic states in the diseased CNS. This has hindered a mechanistic understanding of how distinct phagocyte populations contribute to either injury or repair^9–11^.

Recent advances in high-dimensional single-cell technologies and fate-mapping approaches have enabled more precise dissection of phagocyte identity and function across disease stages of neuroinflammation. These tools have uncovered substantial heterogeneity within CNS phagocyte populations and revealed dynamic transitions in cellular states during neuroinflammation^10^. Despite this progress, the mechanisms by which microglia and MdCs contribute to the chronic neurodegenerative process remain incompletely understood^9,10^. Proposed drivers include oxidative damage^12,13^, iron accumulation, aberrant synaptic remodeling, impaired clearance of myelin debris, and a failure to support remyelination^14^—yet how these processes are regulated and whether they can be selectively modulated remains unclear.

In this study, we use high-dimensional single-cell immune profiling to map changes in the CNS immune cell landscape over the course of progressive neuroinflammation in mice. This approach allowed us to distinguish resident macrophage populations from infiltrating phagocytes, which were characterized by chronic activation as well as increased lysosomal activity and lipid metabolism. Spatial profiling revealed that this program was instructed by local interactions in the leptomeninges and conserved between murine models and human MS. Together, our findings provide new insights into the cellular and molecular programs of progression, thereby offering potential avenues for therapeutic targeting in progressive MS.

## Main

### Progressive neuroinflammation is characterized by pronounced tissue damage despite low immune infiltration

To map the temporal dynamics of cell state changes in progressive neuroinflammation, we utilized the murine model of experimental autoimmune encephalomyelitis (EAE) in non-obese diabetic (NOD) mice. In these mice, the disease is characterized by a brief acute phase of relapsing-remitting neuroinflammation accompanied by clinical symptoms such as paralysis, which develops into a chronic disease with progressive worsening of clinical and neuropathological features (**Figure 1A**), resembling some aspects of human SPMS^15–18^. Here, we analyzed the CNS of mice in different stages of EAE: the homeostatic CNS (day 0, control), clinical relapse (day 14, acute) and remission (day 21, remission), as well as chronic disease (day 30, chronic) and progression (day 60, progressive). Despite similar clinical scores compared to chronic disease, we selected day 60 as the “progressive” time point to capture the underlying mechanisms that cause, rather than associate with progression (**Figure S1A)**.

**Figure 1.**
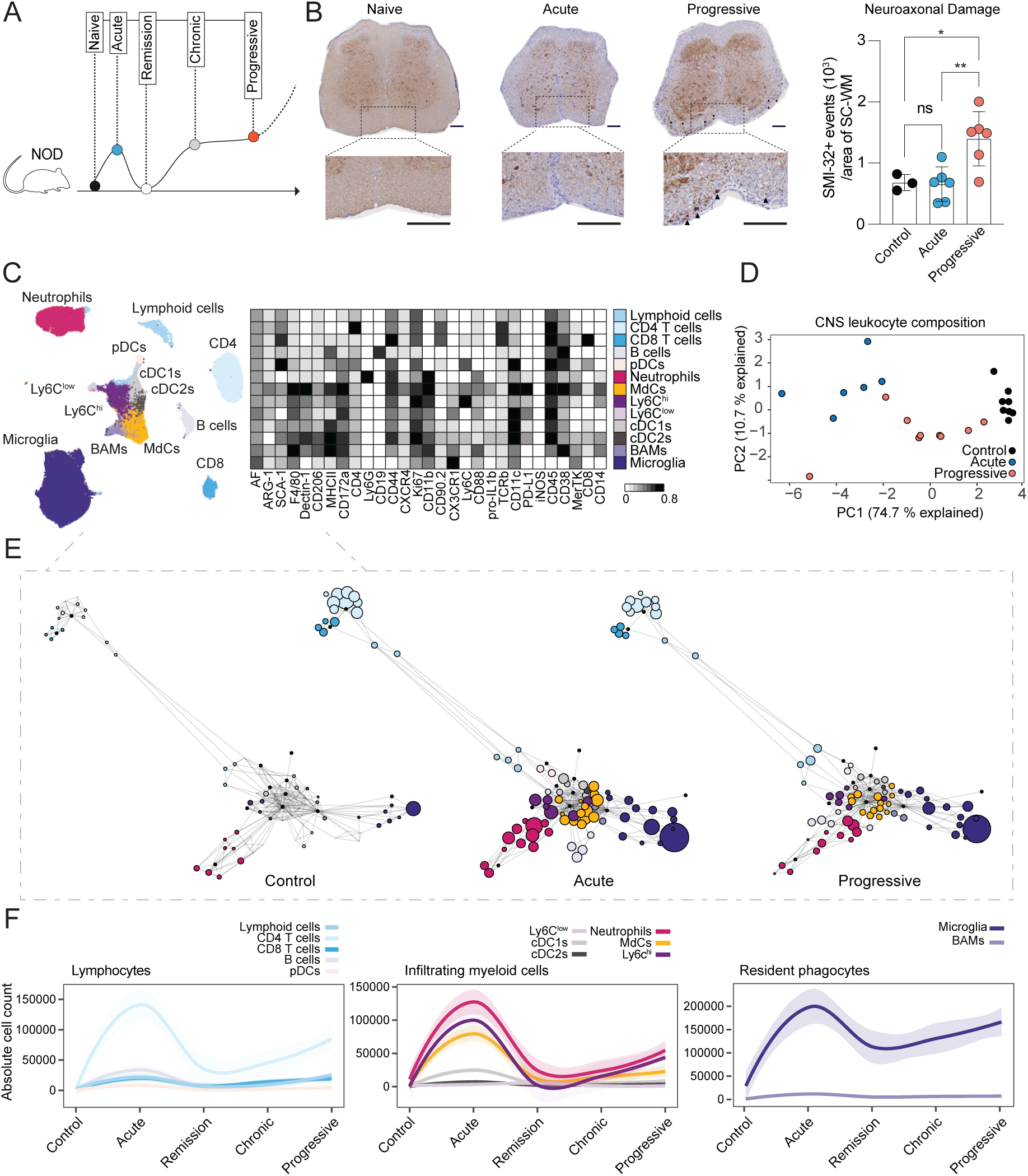
Progressive neuroinflammation is characterized by pronounced axonal damage and reduced immune infiltration. (**A**) Schematic illustration of the clinical score in the non-obsese diabetic (NOD) EAE murine model of progressive neuroinflammation. Each dot depicts a disease stage: control (black), acute (blue), remission (white), chronic (grey), and progressive (red). (**B**) Left: Representative non-phosphorylated neurofilament (nNF) SMI-32 staining in lumbar spinal cord (SC) of control, acute and progressive EAE. Scale bar of 100 uM. Right: The average count of SMI-32^+^ elements detected per white area from lumbar SC sections in control mice (unimmunized, n=3, m/f) and mice with acute (n=6, m/f) and progressive (n=6, m/f) EAE (One-way ANOVA and Tukey’s multiple comparison; control vs progressive *P = 0.0298; acute vs progressive **P = 0.0072). Data pooled from two independent experiments. (**C**) UMAP (left) and heatmap (right) of CNS leukocytes detected in the CNS of NOD mice. UMAP displaying the leukocyte composition in the CNS of randomly subsampled immune cells from all combined disease stages together. Color coding indicates cell type determined by FlowSOM clustering and manual merging. Heatmap displaying the normalized expression profiles of immune clusters annotated correspond to resident macrophages (microglia and BAMs), infiltrating myeloid cells (monocyte-derived cells: MdCs, Ly6c^hi^ and ^low^ monocytes, neutrophils, conventional dendritic cells 1 (cDC1s) and 2 (cDC2s)), lymphocytes (CD4^+^ and CD8^+^ T cells, B cells and other lymphoid cells) and plasmacytoid dendritic cells (pDCs). (**D**) Principal component analysis (PCA) of CNS leukocyte composition based on absolute cell counts for leukocyte clusters indicated above. Dots represent individual mice and are colored according to disease stage. (**E**) Force-directed layouts displaying 100 cell clusters colored by CNS leukocyte annotation in (**C**) and split into control, acute, and progressive disease stages. (**F**) Pseudo time-course displaying the smoothed conditional mean of absolute cell counts of lymphocytes, infiltrating myeloid cells and resident phagocytes obtained by non-linear regression analysis. Shaded area represents the 95% confidence intervals. (**C-F**) Data shown corresponds to: control (CFA-immunized, n=8, m/f), acute (n=6, m/f) remission (n=7, m/f), chronic (n=8, m/f) and progressive (n=9, m/f) EAE. Data from one out of two independent experiments.

Unlike human studies, the staggered immunization protocol facilitated the simultaneous analysis of tissue damage and the characterization of the CNS immune cell landscape, allowing for unambiguous distinction between resident immune cells and infiltrating leukocytes. Histopathological analysis confirmed pronounced demyelination in the spinal cord (SC) of mice in progressive disease compared to control mice (**Figure S1B-C).** Staining with Iba1, a pan-myeloid cell marker revealed higher Iba1-immunoreactive area in the SC of mice in acute disease compared to control mice (**Figure S1D-E**). However, there was a trend towards lower Iba1^+^ area in the SC of mice in progressive compared to acute disease, indicative of reduced inflammation (**Figure S1D-E**). In line with human MS^19,20^, serum levels of neurofilament (NfL) and glial fibrillary acidic protein (GFAP), markers of axonal damage and astrocytic activation, respectively, were significantly elevated during both the acute and progressive disease compared to controls, with GFAP levels tending to be higher in the progressive phase **(Figure S1F**). Notably, we observed increased neuroaxonal damage (SMI-32^+^ cells within the white matter) (**Figure 1B-C)** in the SC of mice with progressive disease compared to those with acute disease, similar to SPMS in humans^21,22^.

Next, we performed full-spectrum flow cytometry to interrogate immune cell composition changes across the individual stages of the disease. Unsupervised clustering yielded the major resident and infiltrating leukocyte subsets in the CNS allowing us to trace the abundance of individual populations across disease stages (**Figure 1C-F**). Principal component analysis (PCA) of the global immune composition revealed that the immune cell landscape during the progressive phase was distinct from the acute phase of neuroinflammation. It resembled an intermediate stage of immune cell abundances observed during acute inflammation and the composition in control mice (**Figure 1D-E**). Dissecting individual population dynamics revealed that most CNS-infiltrating leukocytes accumulated during the acute phase, decreased during the resolution of inflammation, and rebounded in a secondary increase during the chronic and progressive stages of the disease (**Figure 1F** and **S1G**). The secondary infiltration during the progressive phase of the disease, however, demonstrated drastically reduced numbers of cellular infiltrates compared to the acute phase (**Figure S1G**). Of note, also resident macrophages (i.e. microglia and border-associated macrophages (BAMs)) increased in numbers during acute, chronic and progressive phases indicating local proliferation in response to inflammation and/or CNS injury^9,23^.

In conclusion, time-resolved immune profiling of CNS leukocytes across progressive neuroinflammation in the NOD EAE model revealed a secondary but less pronounced immune cell infiltration pattern in progressive neuroinflammation. Alongside its histopathological features, the CNS inflammation in the progressive phase of the NOD EAE model reflects human MS pathology, which is characterized by a compartmentalized inflammatory response in the absence of overt blood-brain barrier leakage^6^. Accordingly, immune cell abundances in the CNS alone do not explain the clinical transition from a relapsing-remitting to a progressive disease course.

### Progressive neuroinflammation is characterized by a chronic monocyte-to-phagocyte differentiation

To uncover cellular and molecular drivers of progressive neuroinflammation, we next interrogated cell state deviations of the major leukocyte subsets across the course of progressive EAE in NOD mice. PCA of the median protein expression across all detected CNS-resident and -infiltrating leukocyte populations yielded a clear distinction between mice in acute and progressive neuroinflammation and controls (**Figure 2A left panel** and **S2A).** Unlike the analysis of immune cell abundance, unsupervised analysis of cell state changes indicated that disease-associated phenotypes were not only maintained but surpassed during progressive neuroinflammation, with higher PC1 values observed compared to mice during acute neuroinflammation, relative to controls (**Figure 2A left panel** and **S2A**). This observation contrasted with the PCA of the immune cell composition, indicating that chronic cellular states rather than cell type abundances could be the immunological driver for disease progression. PC1 explained more than 50% of the total variance and recapitulated the clinical disease score across stages of inflammation, suggesting that PC1 may represent a chronification axis, distinguishing chronic and progressive disease stages from acute relapse (**Figure 2A right panel**). When assessing the relative contribution of individual subsets to this inflammatory axis, we observed that MdCs (comprising monocyte-derived macrophages and monocyte-derived dendritic cells^24^) and conventional dendritic cell type 2 (cDC2) phenotypes together accounted for almost 50% of the variance, followed by neutrophils and Ly6C^hi^ monocytes (11% each; **Figure 2B**). This analysis suggested that infiltrating mononuclear phagocytes actively or passively, contribute to progressive neuroinflammation. When analyzing the relative contribution of functional markers to PC1, we found an overrepresentation of the immune checkpoint molecule programmed cell death ligand 1 (PD-L1) (**Figure 2C**). Ranking the relative contribution of each marker revealed that PD-L1 expression in cDC2s and MdCs, respectively, accounted for the majority of the top 5 contributors to the variance of PC1 **(Figure 2C**). Moreover, PD-L1 expression in cDC2s and MdCs, but also in Ly6C^hi^ monocytes, microglia, and neutrophils, strongly correlated with clinical disease score, with the progressive stage showing the highest expression of the checkpoint molecule (**Figure 2D** and **S2B**).

**Figure 2.**
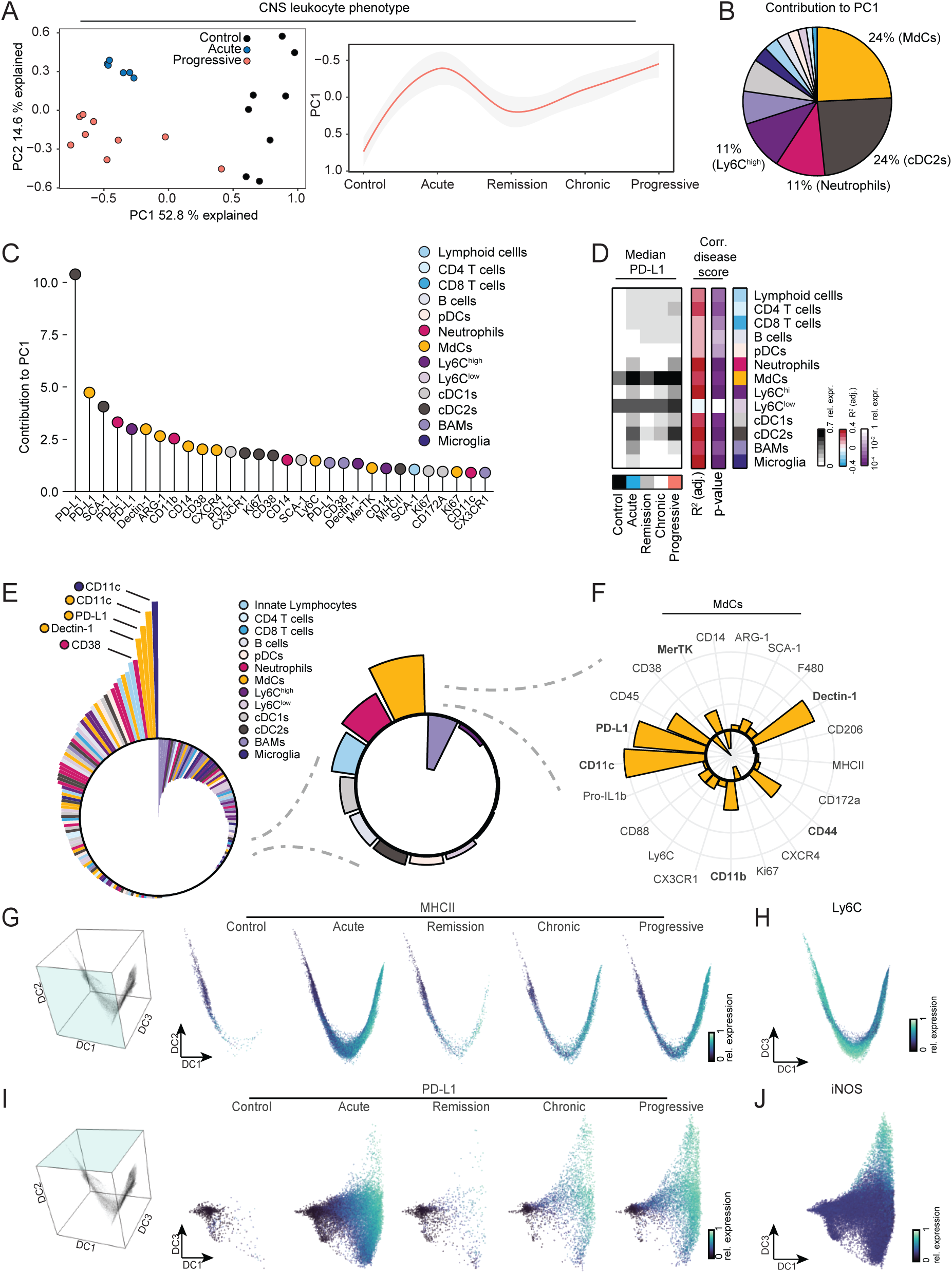
Full-spectrum cytometry profiling reveals chronic monocyte-to-phagocyte transition. (**A–C**) Principal component analysis (PCA) of CNS leukocyte phenotypes based on median marker expression across detected leukocyte subsets. (**A**) PCA of control mice, acute and progressive stages. Dots represent individual mice, and color represents the disease stage (left). Pseudo time-course displaying the smoothed conditional mean of PC1 obtained by non-linear regression analysis across disease stages. Shaded area represents the 95% confidence intervals (right). (**B**) Pie chart displaying the contribution of indicated leukocyte subsets to PC1. (**C**) Lollipop plot depicting the contribution of individual marker expressions of indicated leukocyte subsets to PC1. (**D**) Heatmap showing the median of relative PD-L1 expression across different disease stages for the indicated subsets (left panel). Right color represents R^2^ (adjusted) and purple color p-value for the correlation of median PD-L1 expression with the clinical disease score using a linear model. (**E**) Radar plot displaying the chronification score for indicated median marker expression of the respective leukocyte subsets (left panel), the sum of chronification scores for indicated leukocyte subset (middle panel) and (**F**) the chronification score for the indicated marker expressions in monocyte-derived cells (MdCs). Chronification scores were computed by multiplying the Cohens D effect size of median marker expression between the progressive and acute stage with the Cohens D effect size of median marker expression between the progressive EAE and control mice (**Figure S2D-E**). (**G**) Diffusion maps displaying the diffusion component (DC)2 and DC1, indicative of monocyte-to-phagocyte differentiation (left). Color indicates relative MHCII expression in DC2 vs DC1 across the distinct disease stages (right). (**H**) Diffusion map from all disease stages displaying DC2 and DC1 colored by Ly6C expression. (**I**) Diffusion maps displaying DC1 and DC3, indicative of the chronic phagocyte differentiation (left). Color indicates relative PD-L1 expression in DC3 vs DC1 across the distinct disease stages (right). (**J**) Diffusion map from all disease stages displaying DC3 and DC1 colored by iNOS expression. Data shown corresponds to: control (CFA-immunized, n=8, m/f), acute (n=6, m/f) remission (n=7, m/f), chronic (n=8, m/f) and progressive (n=9, m/f). Data from one out of two independent experiments.

Disease severity and progression are two partially overlapping phenomena that are challenging to untangle. For instance, we found PDL-1 expression in MdCs and cDC2s to strongly correlate with disease severity, but this does not necessarily implicate that PD-L1 expression in MdCs and cDC2s could represent a hallmark of progressive disease (**Figure S2C**). To address this, we developed a pragmatic approach to systematically identify cellular features associated with chronic disease— specifically, markers for which mice in the progressive stage exhibit more extreme values compared to those in the acute stage, relative to control mice. We found that the feature-wise product of Cohen’s d effect sizes—comparing acute to progressive stage and control to progressive stage—satisfied these criteria. Therefore, we refer to the resulting heuristic as “chronification score” (see **Methods** and **Figure S2D**). Accordingly, chronification scores close to zero could be associated with features that do not show major changes between the control, acute, and progressive stages, such as for the expression of F4/80 in pDCs (**Figure S2E**). Negative chronification scores, as assigned for instance to the PD-L1 expression in BAMs, could be interpreted as disease resolution, where protein expression differs from control levels during the acute stage but reverts to resemble control levels in the progressive stage (**Figure S2E**). High chronification scores were assigned to immune features displaying more extreme phenotypes during progressive disease compared to acute (in relation to control mice; **Figure 2E left panel** and **S2E)**. Application of this score to all protein-immune subset combinations, followed by aggregation for individual leukocyte clusters (**Figure 2E right panel**) indicated that cell state alterations in MdCs were most strongly associated with chronic inflammation (**Figure 2F**). When dissecting the contribution of individual markers to the chronification scores in MdCs, we observed high chronification scores for the expression of CD11c, Dectin-1, PD-L1, CD45, CD11b, CD44, and MerTK (**Figure 2F** and **S2F**), resembling some of the disease-associated microglia signatures previously reported in the context of Alzheimer’s disease^25^. The increased expression of MerTK and PD-L1, for instance, was indicative of strong phagocytic activity and immunosuppressive behavior. Notably, chronification scores for lymphocytes or tissue-resident phagocytes, such as microglia and BAMs, were considerably lower, suggesting a relaxation towards a non-inflammatory, homeostatic state during progressive disease.

The phenotypic changes in MdCs associated with chronic stages of neuroinflammation were suggestive of an altered monocyte-to-phagocyte transition during progressive neuroinflammation. To test this hypothesis, we next investigated the differentiation of CNS-infiltrating monocytes. For this, we modeled the monocyte-to-phagocyte differentiation as a continuous process using diffusion maps, a dimensionality reduction method that is particularly well-suited to display cellular differentiation trajectories based on protein expression similarity^24^. Diffusion components 1 and 2 recapitulated the monocyte-to-phagocyte transition characterized by gradual loss of Ly6C expression (**Figure 2H**) and acquisition of *bona fide* macrophage markers such as MHCII, MerTK, F4/80, CD88, and CD14 (**Figure 2G** and **S2G**). Comparison of MdCs across the different stages of neuroinflammation revealed a higher proportion of terminally differentiated macrophages during acute, chronic and progressive stages of the disease, while MdCs in control mice and mice in remission were less differentiated (**Figure 2G**). Strikingly, when analyzing the differentiation trajectory across diffusion component 3, we observed that MdC differentiation in the chronic and progressive stage took a distinct cell fate characterized by high expression of PD-L1 (**Figure 2I**) and Dectin-1 (**Figure S2H**), both of which were assigned a high chronification score before. Moreover, macrophages in the progressive stage exhibited high nitric oxide synthase (NOS)-2 (iNOS) expression (**Figure 2J and S2H**). Recently, iNOS^+^ infiltrating phagocytes have been associated with broad rim lesions predicting rapid progression in people with MS^26^.

In conclusion, our unbiased characterization of major resident and infiltrating immune populations uncovered that chronic monocyte-to-macrophage differentiation is strongly associated with disease progression.

### Monocyte-derived cells during progressive neuroinflammation are characterized by high lipid metabolism and lysosomal activity

To identify the underlying transcriptional circuits and communication networks driving progressive macrophage differentiation, we next performed single-cell RNA sequencing (scRNAseq) of CNS leukocytes at steady state, during acute and progressive EAE, respectively. Graph-based clustering of gene expression profiles identified similar leukocyte subsets as detected using full-spectrum cytometry (**Figure 3A** and **S3A**). Due to the continuous nature of the differentiation trajectory Ly6C^hi^ monocytes and MdCs were analyzed as a joint population in the scRNAseq analysis (termed MdCs). In line with our previous profiling of leukocytes across the stages of neuroinflammation, we found the relative abundance of the major immune subsets to be altered in progressive compared to acute neuroinflammation (**Figure S3B)**. In line with our previous full-spectrum cytometry findings on the protein level, differential abundance analysis revealed that monocytes and their progeny (MdCs) exhibited the highest fraction of altered cellular states in the progressive phase (**Figure 3B-C**), followed by microglia. To pinpoint the molecular differences in MdC states between acute and progressive neuroinflammation, we computed differentially expressed genes of MdCs. MdCs of mice during progressive neuroinflammation were characterized by increased expression of *Fabp5*, *Arg1* (arginase), *Spp1* (osteopontin), *Gpnmb, Ccl6, Cd68, Cd63* and *Plin2,* and a decreased expression of *Ccr2*, *Stat1*, *Ifitm1* and *Ifitm3* compared to MdCs found during acute neuroinflammation (**Figure 3D** and **Supplementary Table 1**). Overall, MdCs in the progressive phase were characterized by reduced NF-κB signaling, and an altered chemokine profile. In contrast, transcripts involved in lipid metabolism, tissue remodeling and lysosomal signaling, were elevated compared to the acute phase (**Figure 3E** and **Supplementary Table 1)**. In addition, we detected increased transcript expression of *Clec7a* (encoding Dectin-1), *Itgax* (CD11c), *Itgam (*CD11b), *Cd274* (PD-L1), *Cd44* and *Mertk* in the progressive stage (**Figure 3F**) that showed high chronification scores in the full-spectrum flow cytometry, further validating our previous findings. Together, our unbiased analysis during progressive neuroinflammation revealed a gene expression program in MdCs skewed towards lipid-metabolism, tissue-remodeling, and lysosomal activity.

**Figure 3.**
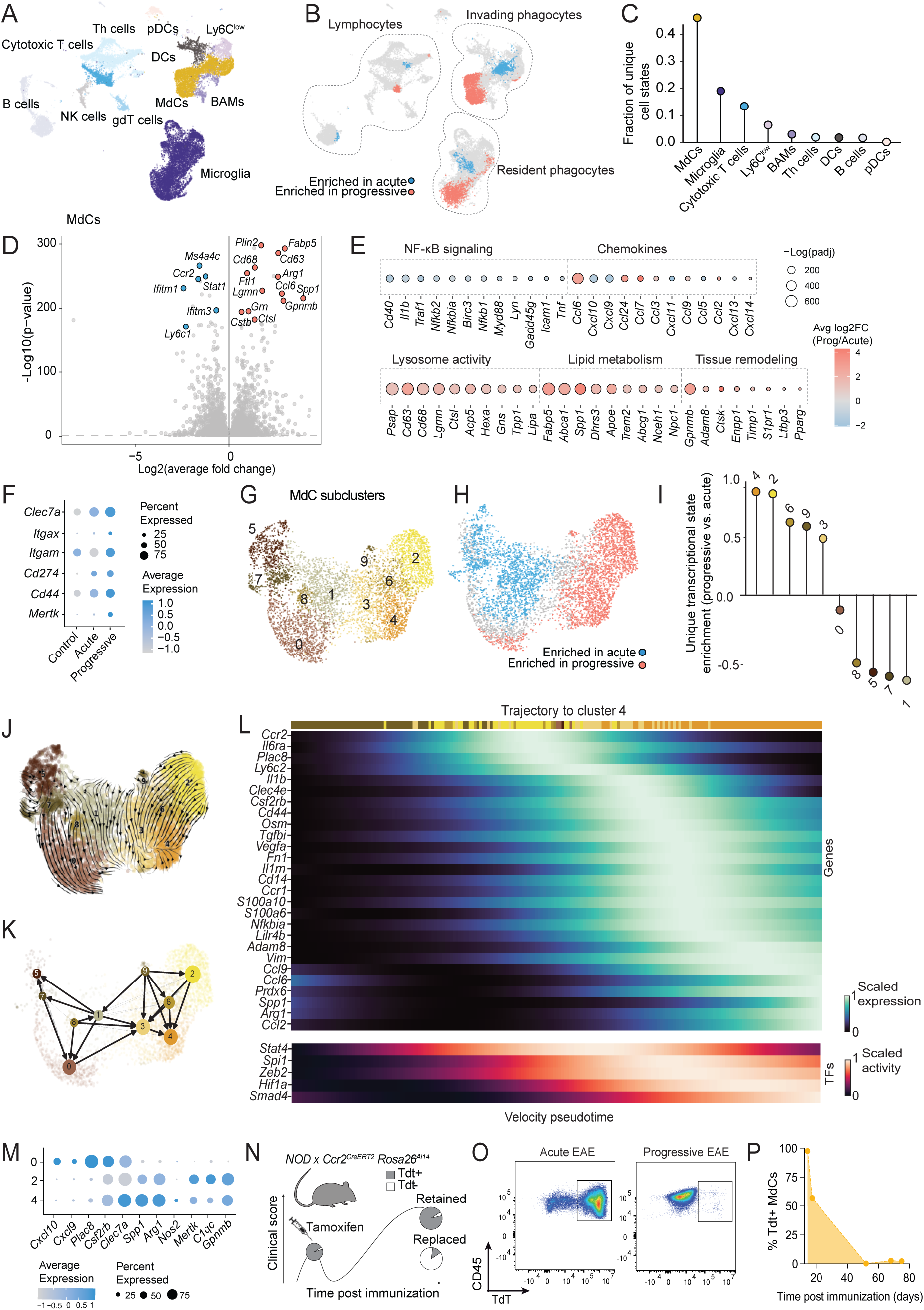
Disease progression is associated with the emergence of lipid-associated macrophages (LAMs) (**A**) UMAP depicting scRNA-seq data of CNS leukocytes in NOD control mice and mice with acute and progressive EAE (**B**) UMAP of the overlay of unique transcriptional states computed by differential abundance scoring on the KNN graph using DA_seq. Blue dots indicate transcriptional states enriched in acute and red dots transcriptional states enriched in progressive EAE. (**C**) Lollipop of the fraction of cellular states enriched in progressive stage grouped by leukocyte subsets. (**D**) Differentially expressed genes (DEGs) in monocyte derived cells (MdCs) when comparing progressive to acute stage of the disease. Blue dots indicate markers upregulated in acute and red dots markers upregulated in progressive EAE. (**E**) Dotplot of the average fold change of gene expression in MdCs of acute and EAE depicting transcripts involved in NfkB signaling, chemokine expression, lysosome activity, lipid metabolism, and tissue remodeling. Circle size is depicting -log 10 of the adjusted p value and color is depicting fold change expression levels. (**F**) Dotplot of markers expression in MdCs in the depicted stages of NOD-EAE. Circle size is depicting % of normalized expression detected per disease stage and color is depicting expression level as average expression. (**G-H**) UMAP of MdCs subclusters (**G**) and of the enriched transcriptional states states (**H**) in acute and progressive disease stages. Blue dots indicate transcriptional states enriched in acute and red dots transcriptional states enriched in progressive. (**I**) Lollipop of the fraction of cell states grouped by MdCs subcluster enriched in progressive stage. (**J**) UMAP displaying the RNA velocity of MdCs. Cells from all disease stages were combined for this analysis. (**K**) PAGA displaying the differentiation probabilities of MdCs in the progressive stage. (**L**) Heatmap displaying smoothed gene expression dynamics (top) or inferred transcription factor regulon activities (bottom) that correlate with the absorption probabilities towards LAM cluster 4. Transcription factor activity has been estimated based on target transcript expression using the decoupler package. (**M**) Dotplot of identity makers expression in MdC subclusters 0, 2 and 4. Circle size is depicting % of normalized expression detected per disease stage and color is depicting expression level as average expression. (**N**) Scheme for *Ccr2*^creERT2/+^ Rosa26^Ai14/+^ x NOD to fate map Ccr2^+^ cells in the CNS. Mice were treated with tamoxifen at day (d)7, d9, d11 and d13. **(O**) Pseudocolor dotplots depicting positive Ai14 (Tdt^+^) MdCs in acute and progressive stages. (**P**) Percentage of Tdt^+^ labelling in MdCs across the course of progressive EAE. For (**A-C**) data shown corresponds to sorted leukocytes (n=27312 cells) from control (unimmunized, n=4401 cells), acute (n=10453 cells) and progressive (n=12458 cells) stages. For (**D-M**) data shown corresponds to subset MdCs (n=5624 cells) from control (n=245 cells), acute (n=2185 cells) and progressive (n=3194 cells). For (**N-P**) data corresponds to: d14 (n=2, m/f); d17 (n=1, f); d52 (n=2, f); d68 (n=1, m); and d75 (n=3, m/f) post immunization.

To interrogate the transcriptional programs associated with the altered monocyte-to-macrophage transition during progressive compared to acute neuroinflammation, we performed an in-depth analysis of differentially abundant cell states in the monocyte compartment. We found that the progressive phase was associated with distinct, transcriptionally defined clusters of MdCs compared to the acute phase (**Figure 3G-H**). Specifically, we identified two clusters of mature MdCs (cluster 2 and 4) that were unique for the progressive stage (**Figure 3H-I**). Their specific transcriptomic profile was characterized by *Apoe*, *C1qa*, *Gpnmb*, *C1qb*, *C1qc* for cluster 2 and *Arg1*, *Cxcl2*, *Clec7a*, *Spp1*, *Ccl6*, and *Ccl24*, for cluster 4, respectively (**Figure S3C**). Given the large overlap of this transcriptional signature with previously described lipid-associated macrophages (LAMs)^27^—characterized as phagocytic cells with lipid-handling capacity across diverse metabolic and inflammatory conditions—we refer to the established term “LAMs” to describe this population. This decision reflects both the conserved features of these cells across disease contexts and tissues, and their apparent independence from ontogeny, supporting conceptual clarity and alignment with emerging consensus in the field^28^.

### Progressive neuroinflammation is characterized by the differentiation of newly infiltrating monocytes towards *bona fide* LAMs

To investigate the differentiation trajectory of infiltrating monocytes towards LAMs, we next computed the ratio of spliced to unspliced transcripts for the monocyte lineage and utilized CellRank to model the cell fate trajectory of LAMs across the course of neuroinflammation (all disease stages combined). We observed that cells early along the differentiation trajectory (cluster 1, 5, 7, 8 and 0) (**Figure 3J**) displayed gene signatures of less differentiated CNS-infiltrating MdCs (*Ccr2, Sell1* and *Ly6a;* **Figure S3C**). These cells transitioned into *bona fide* LAMs (clusters 2 and 4), which were predicted as late stages of the differentiation trajectory (higher velocity pseudotime values, **Figure S3D**).

When modelling the differentiation trajectory of monocytes during progressive neuroinflammation separately, we identified cluster 0 as a potential monocyte precursor of LAMs and cluster 4 and 2 as the terminal stages (**Figure 3K** and **S3E-G**). The transition towards LAM cluster 4 was characterized by a transient expression of inflammatory genes such as *Cd44*, *Ccr1*, *Cd14*, *S100a6*, along gene programs associated with wound-healing (*Tgfbi*, *Fn1, Vegfa*) and the subsequent acquisition of LAM features *Ccl6*, *Arg*1, *Spp1* (**Figure 3L-M**). Interestingly, this differentiation was predicted to be driven by transient *Zeb2* followed by sustained *Hif1a* transcription factor activity (**Figure 3L**). *Zeb2* was recently described as a master regulator of tumor-associated macrophages, suppressing gene programs that control antigen presentation and inflammation^29^. *Hif1a* (hypoxia-inducible factor 1-alpha) is involved in the induction of wound-healing and tissue repair gene programs^30,31^ , and genetic variants in *HIF1A* in people with MS have been recently associated with the risk of a progressive disease course^32^. Conversely, LAM cluster 2 was characterized by a transient increase of LAM signature genes (*Selenop, Gpnmb, Trem2*) along transcripts involved in complement activation (*C1qc, C1qb*) and antigen presentation (*H2-Aa, Cd74,* and inferred CIITA activity; **Figure 3M** and **S3I**). When assessing transcripts encoding proteins that we have previously associated with disease progression using flow cytometry, we found LAM cluster 4 to be characterized by *Clec7a*, *Arg1*, and *Nos2* (encoding iNOS) expressed while *Mertk* was enriched in LAM cluster 2 (**Figure 3M**).

Of note, the potential cellular precursor of both LAM clusters, cluster 0, was characterized by high expression *Cxcl10*, *Cxcl9*, *Tgm2*, *Tgb3*, *Plac8, Irf1* (**Figure 3M** and **S3H**), resembling the recently described *Cxcl10*^+^ monocytes associated with chronic EAE stages^33^. Consistent with previous reports, these cells exhibited high expression of *Csf2rb* (**Figures 3M** and **S3H**), which encodes the receptor for granulocyte-macrophage stimulating factor (GM-CSF). GM-CSF has been previously shown to be involved in pathogenic phagocyte transition in EAE^24,34,35^.

Together, these findings suggest that LAM differentiation is a phenomenon specific for progressive rather than acute disease. To functionally test whether LAMs arise from monocytes recruited *de novo* during the chronic-progressive stage—or whether they represent a persistent population initially seeded during the acute phase—we performed genetic monocyte fate-mapping. For this purpose, we crossed a *Ccr2^CreERT2^ Rosa26^Ai^*^14^ fate mapping strain on the NOD background (**Figure 3N**). Of note, the F1 generation of NOD and C57BL/6 mice still develops progressive EAE^16^. We induced EAE and treated mice with tamoxifen three times (day 8, 10, 12) to achieve Cre-mediated recombination and label all monocytes and their progeny with Tdtomato (Tdt) during the acute phase of the disease (**Figure 3N-O, S3J-K**). This strategy resulted in Tdt labelling of 97.5% of MdCs and 68.15% of T cells, while CNS resident microglia were not labelled at day 14 **(Figure S3K)**. Following MdCs over time revealed a complete loss of Tdt labeling during chronic and progressive disease stages (**Figure 3O-P**), confirming our prediction that monocytes are recruited *de novo*.

In conclusion, we identified a distinct monocyte-to-macrophage differentiation towards *Arg*1^+^ *Spp1*^+^ *Clec7a*^+^ *Nos2*^+^ and *Gpnmb*^+^ *C1qc*^+^ *Mertk*^+^ LAMs as a hallmark of progressive neuroinflammation that may have evolved via a *Cxcl10*^+^, *Csf2rb*^+^ precursor state.

### Local cytokine networks dictate LAM differentiation during progressive neuroinflammation

Our genetic fate-mapping data indicated that LAMs are not retained in the inflamed tissue but are instead replaced by peripheral monocytes over time. Full-spectrum flow cytometry analysis of blood, splenic, and bone marrow monocytes revealed only moderate changes of progressive compared to acute disease in their marker expression, contrary to our observations in the CNS (**Figure S4A**), suggesting that the observed LAM features are imprinted by the chronically inflamed CNS microenvironment.

To identify local tissue cues that may orchestrate the differentiation of LAMs, we next employed Nichenet, a computational method that utilizes knowledge databases linking ligand-receptor interactions with well-defined alterations in downstream targets, to predict cellular interactions^36^. For this, we prioritized ligands expressed in the CNS of mice with progressive disease based on their ability to explain the gene expression dynamics during the differentiation from monocytes to LAMs (**Figure 4A-B**). This analysis indicated a crosstalk between LAMs and CD4^+^ T helper (Th) cells, autocrine and paracrine interactions with MdCs, and interactions with CNS resident microglia as potential drivers of their differentiation (**Figure 4A-B**). Ligands that best explained LAM differentiation included *Csf1* expressed by microglia*, Tgfb1, Il6* and *Il1b* expressed by MdCs, and *Tnf* and *Csf2* expressed by Th cells (**Figure 4B** and **S4B-C**). Of note, *Csf2rb* was predicted as a target for the terminal differentiation towards LAMs (**Figure 3L**). Analysis of the progressive stage of NOD mice, identified Th cells as the major source of *Csf2* (**Figure 4C**). Using full-spectrum cytometry, we confirmed active GM-CSF production by CD4^+^ Th cells and CD8^+^ cytotoxic T cells in progressive disease (**Figure S4D**), in line with previous reports of GM-CSF expression by T cells from MS patients^37^. While a similar fraction of Th cells produced GM-CSF in acute and progressive stages, there was a trend towards increased proportion of cytotoxic T cells producing GM-CSF during progressive disease (**Figure S4E**). Interestingly, analyzing the T cell fraction of the *Ccr2^CreERT2^* fate-mapping experiment revealed substantial Tdt labeling of T cells in progressive stage analysis (**Figure S4F-G**), suggesting that, in contrast to MdCs, a proportion of T cells found during progressive neuroinflammation are retained from the acute stage and continue to produce cytokines.

**Figure 4.**
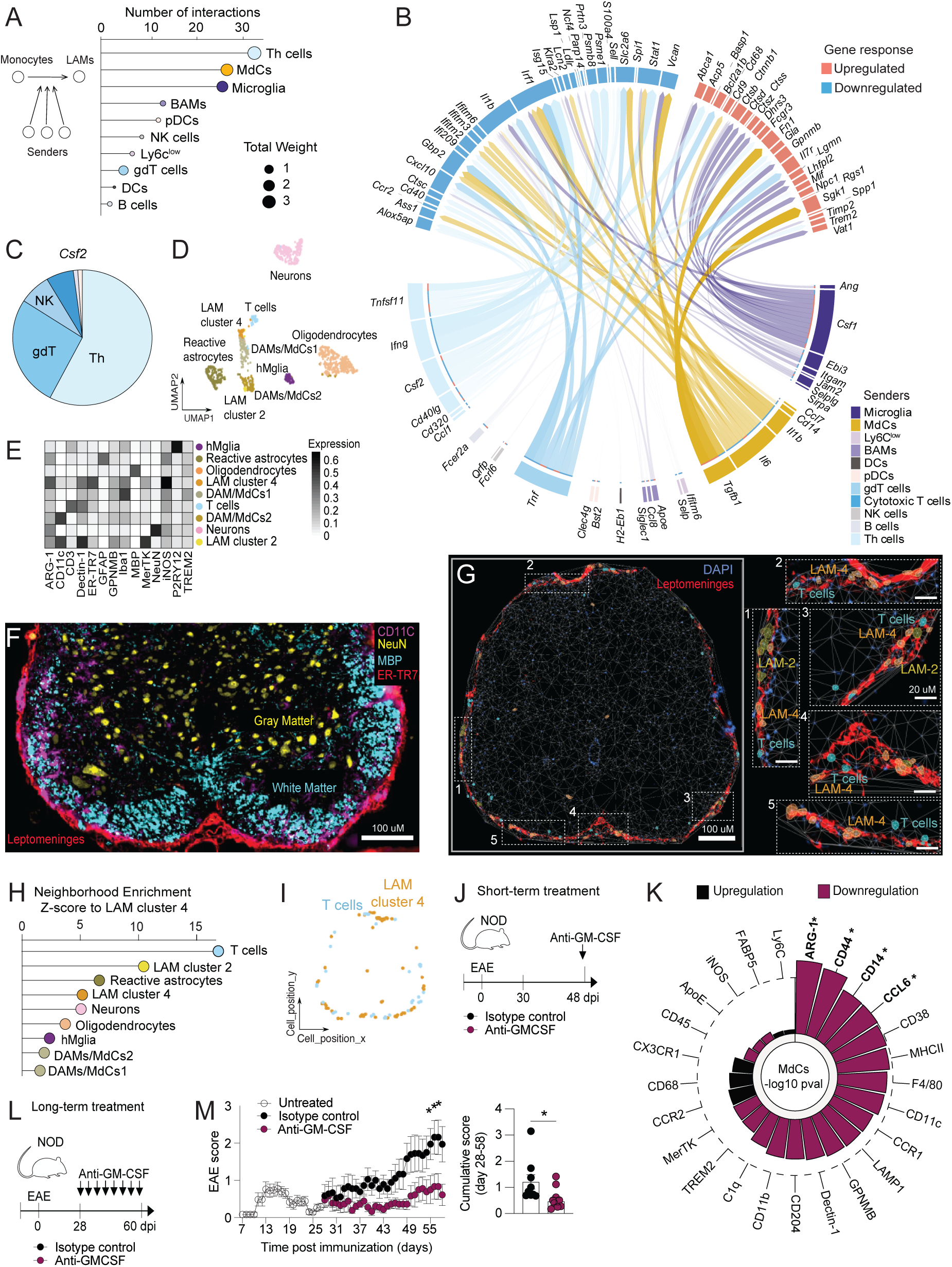
Meningeal T cells promote LAM differentiation via GM-CSF. (**A**) Scheme illustrating Nichenet analysis to predict LAM signature acquisition (left) and lollipop of counts and strength of interactions of sender leukocyte subsets (right). (**B**) Circos plots showing links between unique ligands from senders (ribbon color indicates cluster of origin for each ligand) and predicted associated DEGs in MdCs. Upregulated DEGs are shown in red, downregulated DEGs are shown in blue. Transparency indicates interaction strength, and the ribbon thickness is proportional to the ligand’s regulatory potential. (**C**) Pie chart depicting the distribution of *Csf2* expressing leukocyte subsets in progressive EAE. **(A-C)** Data correspond to sorted leukocytes (n=27312 cells). (**D)** UMAP and (E) heatmap of cell subsets identified using Lunaphore COMET^TM^ multiplex microscopy platform in lumbar progressive spinal cord (SC) sections (progressive, n=4 m/f). Cell subsets correspond to homeostatic microglia (hMglia), reactive astrocytes, oligodendrocytes, LAM cluster 4, (disease-associated microglia) DAM/MdCs 1, T cells, DAM/MdCs 2, neurons and LAM cluster 2. (**F**) Representative lumbar SC section: Red depicts leptomeninges (ER-TR7), magenta depicts white matter (MBP) and yellow depicts gray matter region (NeuN). (**G**) Representative SC overlay of cell subsets: LAM cluster 4, LAM cluster 2 and T cells together with DAPI and ER-TR7 (leptomeningeal fibroblast staining). (**H**) Lollipop of neighbour enrichment Zscore to LAM4. (**I**) Overlay of LAM cluster 4 and T cell distribution across x and y axis. (**J**) Scheme for NOD short-term treatment with anti-GM-CSF (300 μg, three consecutive days) and isotype control during the chronic disease (day 45-48). (**K**) Radar plot depicting the log10 p values of marker differences between anti-GM-CSF treated and isotype control mice. In case a marker was measured in two independent experiments, data from both batches were combined by fitting a linear regression model adding the experiment covariate as a blocking variable. Otherwise, markers were compared using a t test. Bar color denotes the upregulation or downregulation of the respective marker. Dashed line denotes -log10 (0.05) cutoff for p value. (**L**) Scheme for NOD long-term treatment with anti-GM-CSF (300 μg, three times per week) and isotype control during the chronic-progressive stage of the disease (day 28-60). (**M-N**) Plots show mean clinical score over time (two-way ANOVA, and Sidak’s post hoc test, day 55 *P=0.0488, day 56 *P=0.0247, day 57 *P=0.0247, day 58 *P=0.0175) (left) and cumulative clinical score in isotype control treated (n=9, m/f) and anti-GM-CSF treated (n=9, m/f) mice. Unpaired two-tailed t-test (*P=0.0265) (right).

### LAMs are located in close proximity to T cells in the meninges of the spinal cord

Notably, the predicted crosstalk of T cells and MdCs via *Csf2* is in line with our observation of a GM-CSF-sensing precursor for LAMs. As colocalization in the same or adjacent niches is a prerequisite of cell-cell interactions, we next asked whether LAMs could be found in close proximity to T cells in progressive neuroinflammation. To address this, we performed multiplex immunofluorescence (IF) imaging using the Lunaphore COMET^TM^ platform on lumbar spinal cords from NOD mice with progressive EAE. Leveraging this high-parameter imaging approach combined with single-cell segmentation and high-dimensional clustering, we identified 9 immune and non-immune cell clusters (**Figure 4D-E**), with distinct tissue locations (**Figure S4H**). Of note, we identified two clusters expressing the LAM signatures: LAM cluster 4 (ARG-1, Dectin-1, iNOS) and LAM cluster 2 (GPNMB, MerTK) (**Figure 4E**). These two cell types were found within the SC leptomeningeal region, which exhibits a high immunoreactivity for the fibroblast ER-TR7 marker (**Figure 4F-G** and **S4I**). To investigate the spatial relationships of LAM cluster 4 (the terminal LAM cluster), we calculated a spatial Z-score, which measures how close cell types are located compared to random distribution. This analysis revealed that T cells were much closer to LAM cluster 4 than expected by chance, followed by LAM cluster 2 (**Figure 4H**). Visualizing the location of T cells and LAM cluster 4 in the tissue confirmed that this interaction was likely occurring in SC leptomeninges (**Figure 4I**). Renalyzing a recent spatial transcriptomic data set from chronic C57BL/6 EAE SC lesions by Kukanja *et al.*, revealed a LAM-like gene signature with a similar spatial distribution (**Figure S4J-P**), suggesting leptomeningeal LAM-T cell interactions as a conserved feature of chronic-progressive neuroinflammation in mice.

### Blocking GM-CSF signalling impairs LAM differentiation and ameliorates disease progression

To investigate the functional role of LAMs during progressive neuroinflammation, we next sought to interfere with their differentiation and investigate whether this affected disease progression. Based on the high *Csf2rb* expression of the predicted LAM-precursor, the ability of *Csf2* to explain transcriptional dynamics during LAM differentiation, and the close proximity of LAM cluster 4 to T cells, we next assessed whether therapeutic intervention using GM-CSF blockade affected LAM differentiation. To functionally test this, we treated mice with a GM-CSF blocking antibody or an isotype control (IgG). To avoid interfering with the clinical course of EAE - and to specifically isolate the effect of anti-GM-CSF treatment on MdCs (and potentially other *Csf2rb* expressing cell types; **Figure S5A**), we performed a short-term treatment regimen with daily injections between days 48 and 51 (**Figure 4J**). As anticipated, short-term anti-GM-CSF did not affect the clinical score (**Figure S5B**). Instead, we observed decreased expression of several key markers that were associated with LAM differentiation, including ARG-1, CD44, CD14 and CCL6 (**Figure 4K** and **S5C**). Next, we asked whether functional perturbation of LAM differentiation affected disease progression in mice. Therefore, we performed a long-term treatment using an anti-GM-CSF antibody shortly after remission during the early chronic stages (day 28) (**Figure 4M-N**). Intriguingly, this prevented disease progression, supporting a pathogenic role of LAMs during progressive neuroinflammation (**Figure 4M**).

In conclusion, we demonstrate that GM-CSF signaling is crucial for LAM differentiation. Impaired LAM differentiation was associated with improved clinical scores during progressive neuroinflammation, suggesting that these cells contribute to tissue damage in later disease stages.

### CSF1-producing DAM2-like microglia ameliorate disease progression

Apart from infiltrating monocytes, our systematic profiling of leukocytes across the stages of neuroinflammation revealed that CNS-resident microglia change their phenotype during progressive disease (**Figure 2B-C** and **3B-C**). Analysing differentially expressed genes between microglia from progressive compared to acute disease revealed decrease of transcripts involved in inflammation (*Ccl2*, *Ccl12*, *Tspo*, *Cd74*) and an increase in homeostatic (*P2yr12, Hexb, Cx3cr1*) and disease-associated genes (*Cst3, Cst9, Cd9, Trem2*) (**Figure S5D** and **Supplementary Table 2**). Further dissecting the cell states within the microglia compartment yielded four transcriptionally defined clusters (**Figure 5A** and **S5E-G**). Interestingly, one of the clusters was enriched in DAM2 signature genes (*Ccl6, Cst7, Cd9, Lpl, Axl, Spp1, Cybb, Clec7a* and *Timp2*)^38^ and the abundance of this cluster was higher in the progressive stage of EAE (**Figure 5B-C**). DAM2 microglia have been previously described as the late stage of a disease-associated activation trajectory and have been observed across multiple neurodegenerative diseases (reviewed in^38^). Notably, this cluster was characterized by the high expression of *Csf1* and *Itgax* (**Figure 5D**). Intriguingly, microglial CD11c (encoded by *Itgax*) showed the highest chronification score during our full-spectrum flow cytometry analysis (**Figure 5E** and **S5H**), suggesting persistent activation and confirming DAM2 program acquisition in the progressive disease stage. In line with this, our ligand-receptor inference analysis identified *Csf1* as a key ligand produced by microglia in driving LAM differentiation (**Figure 4B**).

**Figure 5:**
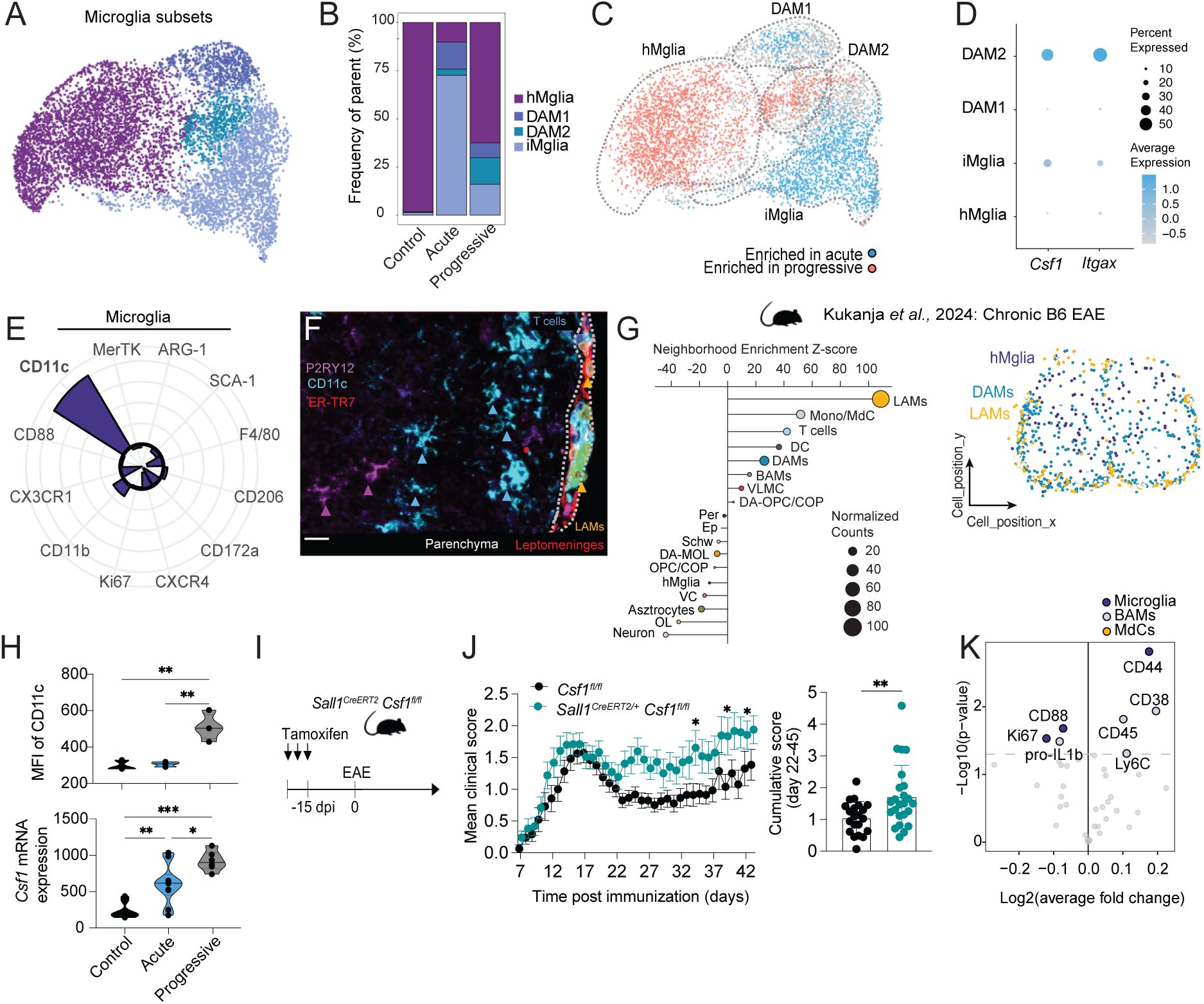
CSF1-producing DAM2-like microglia counteract disease progression in chronic neuroinflammation. (**A**) UMAP of microglia subclusters in NOD mice analyzed by scRNAseq (control, acute and progressive EAE). (**B**) Bar plot of identified microglia subclusters: DAM1, DAM2, iMglia (inflammatory microglia), and hMglia (homeostatic microglia) in the depicted disease stages. (**C**) UMAP of the overlay of unique transcriptional states computed by differential abundance scoring on the KNN graph using DA_seq. Blue dots indicate transcriptional states enriched in acute and red dots transcriptional states enriched in progressive EAE. Dashed lines highlight microglia subclusters. (**D**) Dot plot of average transcript expression of *Csf1* and *Itgax* (encoding CD11c) across the microglia subclusters. (**E**) Radar plot displaying the chronification score for indicated median marker expression in microglia during the progressive stage determined by flow cytometry. (**F**) Representative lumbar SC multiplex immunofluorescence staining of hMglia and DAMs, T cells and LAMs. P2ry12 in purple, CD11c in magenta and ER-TR7 (meninges) in red using Lunaphore COMET^TM^. Overlay of T cells in blue, and of LAMs in yellow (n=4 lumbar SC from progressive EAE). (**G**) Lollipop of neighbour enrichment Zscore to LAMs in chronic C57BL/6 EAE; reanalysed from Kukanja *et al.*^39^(left). Overlay of DAMs (in magenta) and LAMs (in yellow) in lumbar SC (right). Clusters annotated correspond to: LAMs (cluster 3, **Figure S4**), Mono/MdC (monocytes), T cells, dendritic cells (DC), disease-associated microglia (DAMs), border associated macrophages (BAMs), vascular leptomeningeal cells (VLMC), disease-associated oligodendrocyte precursor cells and differentiation-committed precursor (DA-OPC/CO, Pericytes (Per), Ependymal cells (Ep), Schwann cells (Schw), DA-mature oligodendrocytes (DA-MOL), OPC/COP, homeostatic microglia (hMglia), vascular cells (VC), astrocytes, OL (oligodendrocytes), and neurons (further annotated in **Figure S4**). (**H**) Active EAE was induced in C57BL/6 mice by MOG_35-55_/CFA/PT immunization. Upper panel: Median of CD11c protein expression (flow cytometry) and lower panel: *Csf1* mRNA expression (qPCR) in microglia from control (unimmunized), acute (day 15) and chronic EAE (day 24-40). For CD11c expression analysis data correspond to control (unimmunized, n=3, m/f), acute (n=3, m/f), chronic (n=3, m/f) EAE (Kruskall-Wallis ANOVA and Dunn’s multiple comparison; control vs chronic **P=0.0034, acute vs chronic **P = 0.0055). For *Csf1* expression data corresponds to: control (unimmunized, n=4, m/f), acute (n=6, m/f), chronic (n=5, m/f) (Kruskall-Wallis ANOVA and Dunn’s multiple comparison; control vs acute **P=0.0029, control vs chronic ***P < 0.001 and acute vs chronic *P = 0.035). (**I**) Schematic representation of tamoxifen treatment strategy to target *Csf1* in microglia followed by EAE induction (see above). (**J**) Plots show mean clinical score over time (two-way ANOVA, and Sidak’s post hoc test, day 34 *P=0.0457, day 39 *P=0.0469, day 41 *P=0.0356) (left) and cumulative clinical score in *Csf1*^fl/fl^ (n=21, m/f) and *Sall1^CreERT2/+^ Csf1*^fl/fl^ (n=24, m/f) mice. Unpaired two-tailed t-test (**P=0.0094) (right). (**K**) Volcano plot of marker fold change in MdCs, microglia and BAMs. Color dots represent significant changes in the phagocyte subsets. For (**A-D**) data shown corresponds to microglia (n=7712 cells) from control (unimmunized, n=1219 cells), acute (n=2760 cells) and progressive (n=3733 cells) stages. For (**E**) data corresponds to control (CFA-immunized, n=8, m/f), acute (n=6, m/f) progressive (n=9, m/f). All data are shown as mean ± SEM.

Spatial characterization using the Lunaphore COMET^TM^ system revealed that CD11c⁺ microglia preferentially localized near meningeal LAMs and displayed a striking spatial gradient: CD11c expression increased, and cellular morphology became less ramified as cells approached the leptomeningeal space (**Figure 5F**). This anatomical alignment points to a localized crosstalk between the two cell types—where DAM2-like microglia may provide local instructive cues for LAM differentiation through CSF1 signaling within the meningeal niche. Indeed, our findings align with observations by Kukanja *et al.*, that also predicted CSF1-mediated interactions between DAMs and MdCs in late-stage EAE^39^. Similar to progressive neuroinflammation, DAMs in chronic C57BL/6 EAE were located in close proximity to LAMs and T cells in the leptomeninges (**Figure 5G**) and expressed high CD11c (flow cytometry) and *Csf1* (mRNA) (**Figure 5H**). Together, these results support the emergence of LAMs and DAM2 microglia as conserved phagocyte cell states across different models of chronic/progressive neuroinflammation.

CSF1 produced by microglia (**Figure S5I**) in chronic and progressive EAE stages can be sensed by MdCs as predicted in our ligand receptor inference (**Figure 4B**), by CSF1 receptor expressing BAMs and microglia themselves, and to a lesser extent by other cell types of the hematopoietic and non-hematopoietic system **(Figure S5J-K)**. To directly test the functional role of microglial CSF1 production in disease progression, we used a conditional knockout strategy in the chronic C57BL/6 EAE model. This allowed us to circumvent potential off-target effects on peripheral macrophages, as reported in studies using systemic CSF1R antagonism, which have yielded conflicting results^40–43^. Mice harbouring a floxed *Csf1* allele and tamoxifen-inducible Cre recombinase under the *Sall1* promoter were administered three cycles of tamoxifen prior to EAE induction (**Figure 5I**). Two weeks after the final tamoxifen dose, efficient recombination and selective deletion of *Csf1* in microglia was confirmed by qPCR analysis of FACS-sorted cell populations (**Figure S5L**). Previous reports indicated that *Sall1* driven recombination could also target astrocytes^44^. However, our data confirmed that *Csf1* expression was almost restricted to microglia and successfully ablated in Cre^+^ mice (**Figure S5L**). We next induced EAE and monitored motor deficits over time. Strikingly, mice lacking microglial CSF1 failed to recover from paralysis after the initial disease peak and neurological deficits persisted during the remaining course of chronic EAE (**Figure 5J**). These observations suggest a critical role for microglia-derived CSF1 in mediating reparative or neuroprotective processes during disease progression.

Full-spectrum cytometry profiling (**Figure 5K** and **S5M-N**) of CNS myeloid (CSF1R-sensing) subsets revealed an upregulation of F4/80, CD38, and CD45 in BAMs from Cre^+^ mice, consistent with a more activated state, while microglia exhibited reduced Ki67 expression, indicating impaired proliferation in the absence of autocrine CSF1 signalling. Interestingly, we also noted elevated CD44 expression in microglia, a marker previously linked to glial activation and fibrotic scar formation^45^. In contrast, we detected no significant changes in the surface marker phenotype of MdCs between Cre^+^ mice and tamoxifen-treated littermate controls.

Taken together, these findings demonstrate that CSF1-producing DAM2-like microglia are required for tissue homeostasis and functional recovery during chronic neuroinflammation, likely through autocrine survival signalling and modulation of the local microenvironment. Opposed to LAMs derived from infiltrating monocytes, loss of CSF1-producing DAM2-like microglia exacerbated disease severity, suggesting a protective axis of microglial CSF1 in progressive CNS inflammation. Thereby, our systematic single cell profiling of myeloid cells during progressive neuroinflammation in conjunction with genetic ablation models revealed a divergent influence on disease progression for resident and infiltrating phagocytes.

### LAMs are enriched in the meninges of human SPMS patients

To interrogate whether the myeloid cell states associated with progressive neuroinflammation could be found in human disease, we next compared the myeloid landscape of progressive EAE with two publicly available sequencing datasets from human MS patients. Re-clustering of myeloid cells found in Lerma-Martin *et al*.^46^, revealed the presence of three phagocyte clusters: DAMs (enriched for *CTSD, LPL, TREM2, CD9*), homeostatic microglia (hMglia, enriched for *CX3CR*1 and *P2RY12*), and MdCs (enriched in *CD14, CLEC12A* and *ITGA4*) in subcortical MS tissues (**Figure 6 A-B** and **S6A**). Within these phagocyte subsets, we detected the highest LAM signature score within the MdC cluster (**Figure 6C**). When comparing the relative abundance of these LAM-like MdCs, we found that they were almost exclusively detected in the brain of SPMS patients compared to controls (**Figure 6D**).

**Figure 6:**
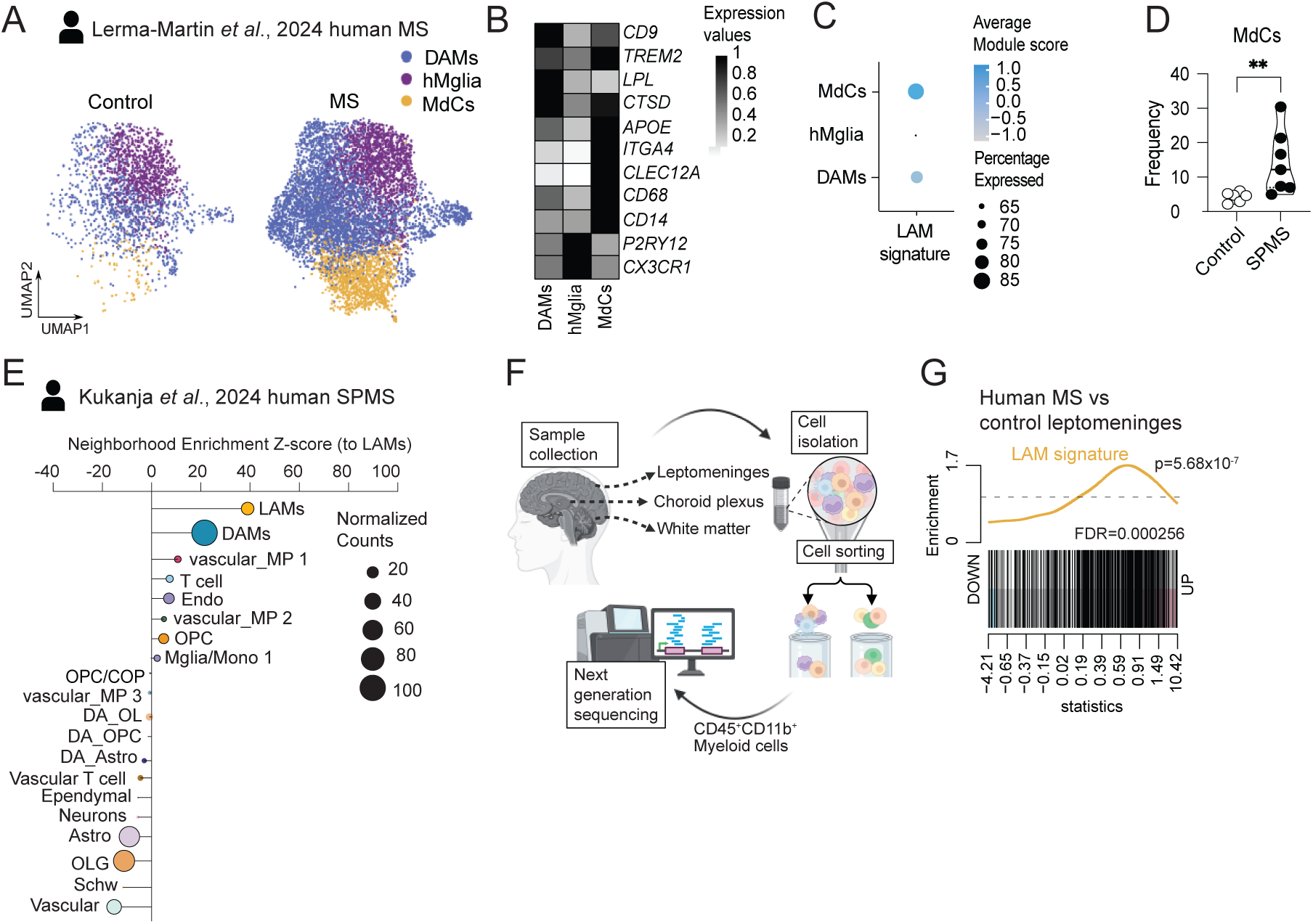
Human SPMS analysis confirms LAM signature and leptomeningeal crosstalk. (**A**) UMAP and (**B**) heatmap of phagocyte subsets in human MS and control subcortical brain tissues reanalyzed from in Lerma-Martin *et al*^46^ (original annotation). (**C**) Dot plot of average module score of LAM signature across the detected phagocyte subsets. (**D**) Frequency of MdCs found in control and MS patients (Mann-Whitney test, **P= 0.0047, data from control (n=6) and MS patients (n=7)). (**E**) Lollipop of neighbour enrichment Z-score to LAMs in human SPMS cervical spinal cord tissue (active lesions) reanalyzed from Kukanja *et al* ^39^. Clusters depicted correspond to LAMs (cluster 15, **Figure S6**), disease-associated microglia (DAMs originally annotated MP/MiGl_2), vascular macrophage (MP) 1, oligodendrocyte precursor cell (OPC), microglia/monocyte 1 originally annotated MP/MiGl_1 (Mglia/Mono 1), oligodendrocyte precursor cells and differentiation-committed precursor (OPC/COP), vascular MP3, disease-associated (DA)-oligodendroglia (DA-OLG), containing annotated clusters (OLG_GM_DA and OLG_WM_DA), DA-OPC, DA-oligodendrocytes (OLs), astrocytes containing Astro_GM_DA and Astro_WM_DA (DA-Astro), vascular T cell, ependymal cells (Ependymal), astrocytes (Astro), OLG (containing OLG_GM and OLG_WM), Schw (schwann cells) and vascular cells (Vascular). (**F**) Schematic representation of experimental strategy to sort and sequence CD11b^+^ CD45^+^ myeloid cells from MS and control from meninges, choroid plexus and white matter. **(G)** GSEA plot based on the LAM signature enriched analysis in myeloid cells from MS vs control patients’ meninges. P value was calculated with the CAMERA gene set permutation test and adjusted for multiple hypothesis testing with the Benjamini–Hochberg method. Data from control (n=7) and MS patients (n=5).

Likewise, analyzing the transcriptome of cell clusters detected within human spinal cord MS active and inactive MS lesions from Kukanja *et al.*^39^, revealed phagocytes that exhibited a LAM-like gene signature (**Figure S6B-E**). When investigating their spatial distribution, we observed that LAMs were located in close proximity to DAMs and vascular phagocytes and, to a lesser extent, in proximity to T cells (**Figure 6E**). Similar to what we observed in chronic/progressive EAE models, human DAMs exhibited high expression of *Itgax,* associated with a DAM2 profile (**Figure S6F**).

Due to the underrepresentation of meningeal tissue in previous studies of human MS, we performed a gene enrichment analysis of bulk RNAseq data from dissected postmortem brain samples of the Netherlands Brain Bank donor program (**Figure 6F**). Intriguingly, this revealed an increase in LAM transcripts in the meninges of MS patients (**Figure 6G**) but not in the normal-appearing white matter or white matter lesions (**Figure S6G**). Notably, there was a trend towards elevated individual transcript expression of ARG-1, a key marker for LAM cluster 4 in meningeal tissue from MS patients compared to controls (**Figure 6H)**. Together, our data reveal a chronic MS progression-associated differentiation trajectory from monocytes to LAMs, characterized by conserved transcriptional features and a shared meningeal niche in both mouse and human.

## Discussion

Despite advances in the treatment of relapsing-remitting multiple sclerosis (RRMS), progressive MS remains therapeutically intractable, largely because the pathogenic mechanisms underlying the transition to chronic neuroinflammation are incompletely understood and are often refractory to systemic immunomodulation. This may be partly explained by the compartmentalized nature of CNS inflammation and the lack of overt immune infiltration during progression^1,6^. Our study addresses critical gaps in understanding the cellular and molecular mechanisms driving chronic neuroinflammation in progressive MS. Using time-resolved single-cell profiling in a murine SPMS model, we identified a critical immunopathological axis centered on MdCs that differentiate into LAMs - a state restricted spatially to leptomeninges and conserved in human SPMS.

An important insight from our study is that immune cell abundance does not necessarily correlate with disease severity - highlighting that severe chronic stages can occur despite reduced immune cell numbers. Despite low inflammatory infiltration, disease progression is marked by distinct molecular profile of MdCs characterized by *Spp1*, *C1qa-c*, *Gpnmb*, *Arg1*, and *Fabp5* expression. These features mirror those described in chronic maladaptive tissue remodeling^27^ across diverse disease contexts, including cancer^47,48^ and neurodegeneration^49,50^, suggesting a conserved, hypoxia-adapted, immunosuppressive state with impaired debris clearance and fibrosis-like features. These wound-healing-like programs may lock the CNS into a state of smoldering damage, particularly at vulnerable sites such as the meninges and cortical rim, where LAMs were most enriched. Notably, the complement cascade—particularly C1q expression by LAMs—emerges as a candidate effector mechanism of damage at the CSF-parenchyma interface ^51–53^, consistent with recent human pathology studies^54–56^. As a component of a broader maladaptive metabolic program, reactive oxygen species (ROS), previously linked to MdC-mediated tissue damage in neuroinflammation^12^, may represent a pathogenic feature of LAMs.

Together, the LAM phenotype—marked by PD-L1 upregulation, lysosomal activation, and lipid handling—may thus represent a maladaptive wound-healing program locked by persistent environmental cues such as GM-CSF and hypoxia. While immune checkpoint molecules often signal inflammatory resolution, depending on disease stage and tissue context, they may reflect an exhausted or suppressive state that fails to resolve tissue pathology. Indeed, impaired PD-1/PD-L1 signaling has been linked to MS progression^57^, and blockade of PD-L1 and other immune checkpoints has shown paradoxical effects in models of CNS autoimmunity and neurodegeneration^58–60^. Whether LAM-expressed PD-L1 modulates local T cell function or acts intrinsically remains to be addressed in future studies^61^.

Our identification of the leptomeninges as a conserved anatomical niche for LAM accumulation aligns with mounting evidence implicating meningeal inflammation in SPMS, especially in driving cortical demyelination and gradient-like neuronal damage^6,62–66^. The presence of LAM signatures in the meninges of MS patients—but not in normal-appearing white matter or lesions—underscores their potential relevance as biomarkers or therapeutic targets. Ligand prediction, spatial transcriptomics and multiplexed imaging reveal physical proximity of LAMs to both *Csf2^+^* Th cells and *Csf1^+^* DAM2-like microglia, supporting a localized cytokine network as a driver of their state.

Mechanistically, we show that progression-associated LAMs are not retained from the acute phase but are continuously replenished from the periphery. Using fate-mapping and transcriptional trajectory analysis, we identify a GM-CSF-sensing, *Cxcl10*⁺ monocyte precursor as the source of these cells. The emergence of *Cxcl10*⁺ monocyte precursors as a transitional population toward LAMs expands upon previous work by Giladi *et al*., who defined *Cxcl10*⁺ monocytes as a pathogenic subset enriched during chronic EAE^33^. While *Cxcl10*^+^ monocytes themselves emerge independently of GM-CSF^24^, our work places these cells upstream of maladaptive macrophage states and implicates GM-CSF as a key differentiation signal towards LAMs. Therapeutically, transient blockade of GM-CSF signaling impairs LAM differentiation and halts progression *in vivo*, positioning GM-CSF as a targetable driver of maladaptive myeloid responses in SPMS. In humans, GM-CSF receptor expression is differentially expressed in monozygotic twins discordant for MS^67^. While prior studies have shown that GM-CSF producing RORγt-positive T cells are required for inducing EAE^68^, they have been described in chronic EAE^69^, as well as in the meninges of SPMS patients^70^. GM-CSF overexpression alone can drive CNS invasion by phagocytes and elicit spontaneous neurologic disease in mice^35^, licensing CCR2⁺ monocytes to adopt pathogenic phenotypes^34^. In addition to those findings, we now describe that in progressive EAE, GM-CSF, primarily derived from meningeal CD4⁺ T cells, drives LAM differentiation and promotes disease progression.

Conversely, microglial production of CSF1 appears to restrict tissue damage by promoting alternative, potentially reparative programs. *Csf1*-expressing microglia adopt a DAM2-like phenotype and exhibit high *Itgax* expression, resembling a reparative program described in neurodegeneration^25^. Conditional deletion of *Csf1* in microglia exacerbates disease in chronic EAE, suggesting a protective role consistent with previous reports showing that CSF1 supports myelin clearance and microglial survival^71^. Finally, our findings help reconcile conflicting data on CSF1R-targeting strategies^42,72,73^. While CSF1R inhibition depletes both protective microglia and pathogenic MdCs, specific loss of microglial CSF1. likely via autocrine signaling^40^ exacerbated chronic disease. This cytokine dichotomy—GM-CSF as a pro-inflammatory differentiation cue versus CSF1 as a protective modulator—mirrors findings in other systems^41^ and offers a framework for interpreting the dual nature of macrophages in neuroinflammation^9,10^. Our results argue for nuanced approaches, potentially leveraging microglia-specific survival and differentiation pathways to counterbalance pathogenic maladaptive phagocyte states.

In summary, our study integrates longitudinal fate mapping, single-cell transcriptomics, and cytokine perturbation to uncover a conserved trajectory of monocyte-to-LAM differentiation as a hallmark of progressive neuroinflammation. By positioning LAMs within a defined cytokine network and anatomical niche, we link our findings with growing evidence implicating compartmentalized inflammation in SPMS, particularly in cortical demyelination and gradient-like neuronal damage. By clarifying the opposing roles of GM-CSF and microglial CSF1, we establish a mechanistic framework for targeting maladaptive phagocyte states while preserving or enhancing reparative microglial functions. These insights position LAMs as central orchestrators of chronic neuroinflammation, and their transcriptional profile, tissue niche, and upstream regulators offer a blueprint for therapeutic intervention in progressive MS. Ultimately, strategies that reprogram LAMs—without disrupting protective microglial circuits—could provide a targeted means to halt or even reverse disease progression in progressive MS.

## Methods

### Mice

All animal care and experiments were performed in adherence with the Swiss federal and institutional guidelines, and all experiments were approved by the Swiss Veterinary Office following 3R guidelines. All mice were bred and maintained by the Laboratory Animals Service Centre (LASC) at the University of Zurich. NOD mice were purchased from Charles River and colonies were maintained in-house. *Ccr2*^creERT2^ were described in^34^ and crossed to Rosa26^Ai14^ described in^74^. *Ccr2*^creERT2/CreERT2^ x *Rosa26^Ai1/Ai^*^14^ mice were further crossed with NOD mice and het F1 were used for reporter experiments. Wild-type C57BL/6 mice were purchased from Janvier Laboratories. *Sall1* ^CreER^ mice were described in^75^ and further crossed to *Csf1*^fl/fl^ described in^76^.

### EAE induction

NOD mice were used as a model of progressive EAE. In brief, active EAE was induced in 12 week-old mice by subcutaneous immunization with 150 μg of MOG_35–55_ emulsified in CFA (4 mg/ml H37Ra strain). *Sall^1CreERT2^ Csf1^fl/fl^* mice (C57BL/6 background) were used for chronic EAE experiments. Here, EAE was induced in 8–10-week-old mice by subcutaneous immunization with 200 μg of MOG_35–55_ emulsified in CFA (1 mg/ml H37Ra strain). For both NOD and C57BL/6 mice pertussis toxin (200 ng) was injected on day 0 and day 2 after immunization.

Clinical signs of EAE were assessed according to the following score: 0, no signs of disease; 1, loss of tone in the tail; 2, hind limb paresis; 3, hind limb paralysis; 4, tetraplegia; 5, moribund.

### In vivo antibody treatment

NOD mice were treated, 300 μg of isotype control antibody (2A3), or 300 μg of anti–GMCSF antibody (anti–GM-CSF; MP1, BioXCell) injected intraperitoneally. Antibody treat schemes are described in the figure legends.

### Tamoxifen treatment

Tamoxifen (Sigma) was dissolved in 2% ethanol and corn oil to 25 mg/ml and administered in 200 μl doses via oral gavage (5 mg per dose).

### Cell Isolation murine samples

Organs were harvested and processed as recently described ^24,77,78^. In brief:

#### Blood

Following CO_2_ euthanasia, peripheral blood was collected via cardiac puncture using heparinized syringes and processed within 2 hours of collection. Red blood cells were lysed using RBC lysis buffer.

#### Bone Marrow

Femurs and tibias were collected prior to perfusion and were cleaned of muscle tissue, and bone marrow was flushed using a 26-gauge needle with ice-cold PBS + 1 mM EDTA. The suspension was filtered through a 70 μm cell strainer and centrifuged at 300×g for 10 minutes at 4°C to obtain a single cell suspension.

#### CNS Leukocytes

Animals were transcardially perfused with ice-cold PBS. CNS tissues (brain and spinal cord) was cut into small pieces and incubated with collagenase IV (0.4 mg/ml) and deoxyribonuclease I (DNase I) (Sigma-Aldrich) for 40 mins and passed through a 19-gauge needle to obtain a single cell suspension. The resulting cell suspensions were enriched by density gradient centrifugation with Percoll (GE Healthcare) (30%) at 2750 rpm for 30 mins at 4°C with no brake, to remove myelin.

#### Spleen

Spleens were harvested and mechanically dissociated using the flat end of a sterile syringe plunger in PBS + 1 mM EDTA. The dissociated tissue was passed through a 70 μm cell strainer and centrifuged at 300×g for 10 minutes at 4°C to obtain single cell suspension.

### Human brain

Cells were sorted from choroid plexus, leptomeninges, normal/lesioned white matter of post-mortem brain from non-neurological control (n=8) and MS donors (n=7) as previously described ^79^. Viable CD45^+^ CD11b^+^ myeloid cells were sorted and analyzed by bulk RNA sequencing ^80^. Reads were aligned to the human GRCh38.p13 genome using STAR (v2.7.10), and gene counts were obtained using HTSeq (v2.0.2). Count data were logCPM-transformed, TMM-normalized (edgeR v4.4.1), and precision-weighted using voom (limma v3.62.2). Differential expression analysis was performed using limma framework. Gene set enrichment analysis was conducted using CAMERA. For each compartment, we performed differential gene expression and gene set enrichment analysis to contrast MS versus control myeloid cells.

### T cell restimulation

Leukocytes of the CNS were restimulated at 37°C in the presence of PMA (50 ng/ml, Applichem) and Ionomycin (500 ng/ml, Invitrogen) for 5 hours, the last two hours in the presence of Golgi Plug and Golgi Stop (1:1000 dilution, BD). Cells were washed three times with ice-cold PBS and subjected to flow cytometry.

### Flow cytometry staining

Prior to surface staining, Fc receptors were blocked with purified anti-mouse CD16/32 (clone 93, BioLegend) for 10 min on ice to prevent non-specific binding. Single-cell suspensions were directly incubated with the surface antibody cocktail for 20 min at 4°C. Cells were washed with PBS buffer and incubated with the fluorochrome-conjugated streptavidin antibodies for 20 min at 4°C. Anti-mouse fluorochrome-conjugated monoclonal antibodies used in this study can be found in **Supplementary Table 3.**

The cells were then washed and fixed for 25 minutes at 4 °C with Foxp3/transcription Factor Staining Buffer Set (Thermo Fisher), or Cytofix/Cytoperm (BD) for cytokines. The cells were stained with antibodies specific for the indicated intracellular epitopes (**Supplementary Table 3**) over night at 4 °C in permeabilization buffer from eBioscience Foxp3/transcription Factor Staining Buffer Set. After washing, cells were analyzed using a Cytek Aurora spectral flow cytometer and analyzed using FlowJo Version 10.8. Cell aggregates and doublets were excluded from analyses using FSC-A versus FSC-H. Dead cells were identified using Zombie NIR Fixable Viability Kit (BioLegend).

### Fluorescence-activated cell sorting for qPCR analysis

Cell sorting of live CD11b^+^CD45^+^CX3CR1^+^CD44^-^ microglia, CNS infiltrating CD11b^+^ CD45^+^ CX3CR1^+^ CD44^+^ Ly6C^+^ Ly6G^-^ MdCs, CD11b^-^ CD45^-^ ACSA-2^+^astrocytes and CD45^-^ ACSA-2^-^ fraction was performed using a using a 100um nozzle on a BD FACS Aria III (FACS DIVA Software v9).

### Quantitative RT–PCR

Total RNA was isolated from sorted microglia, MdCs, astrocytes or CD45^-^ fraction using the QuickRNA Microprep Kit (R1051, Zymo Research) according to the manufacturer’s instructions. Then, complementary DNA (cDNA) was synthesized using the M-MLV Reverse Transcriptase (28025013, Invitrogen) and qRT–PCR was performed on a CFX384 Touch Real-Time PCR Detection System (Bio-Rad) using SYBR Green (Bio-Rad). We calculated gene expression as 2^−ΔCt^ relative to Pol2 as the endogenous control. When comparing cre to wt, relative gene expression was then calculated by dividing cre gene expression value by the average of wt gene expression value.

### Primers for *Csf1*

5′-TACAAGTGGAAGTGGAGGAGCCAT-3′; (forward)

5′-AGTCCTGTGTGCCCAGCATAGAAT-3′; (reverse)

### Primers for *Pol2*

5′-CTG GTC CTT CGA ATC CGC ATC-3′; (forward)

5′-GCT CGA TAC CCT GCA GGG TCA-3′; (reverse)

### Single cell RNA sequencing

Cell Sorting: Live CD11b^+^ CD45^+^ CX3CR1^+^ CD44^-^ microglia, CNS infiltrating CD11b^+^ CD45^+^ CX3CR1^+^ CD44^+^ Ly6C^+^ Ly6G^-^ MdCs and CD11b^-^ CD45^+^ lymphocyte fractions were sorted using a 100um nozzle on a BD FACS Aria III (FACS DIVA Software v9) from control (unimmunized), acute and progressive NOD EAE mice (n=1; clinical score 2.75). Microglia, MdCs and Lymphocytes were combined in equal ratio for each mouse and loaded into 10x Genomics Chromium in parallel. Libraries were prepared as per the manufacturer’s protocol (Chromium Next GEM Single Cell 3ʹ Reagent kits v3.1 protocol) and sequenced on an Illumina NovaSeq sequencer according to 10x Genomics recommendations (paired end reads, R1 = 28, i7 = 8, R2 = 91) to a depth of around 50,000 reads per cell. Initial processing was done using CellRanger (v6.0.2) mkfastq and count. Reads were aligned to the *Mus musculus* reference genome GRCm39, annotated with GENCODE M26 (release 2021-04-20). Data were analyzed using *Seurat v4* ^81^. Ambient RNA contamination was removed using *SoupX* (default parameters). Quality control was performed by filtering out cells with >8% mitochondrial RNA content, fewer than 350, or more than 8,000 detected genes. The remaining cells were normalized to the library size, log1p-transformed, and scaled.

Highly variable genes were identified using the vst method in *Seurat*, selecting the top 2,000 HVGs for downstream analysis. PCA was performed, and the top 50 principal components were used to construct a shared nearest neighbour graph, which was clustered using the Louvain algorithm and visualized via UMAP (all *Seurat*, default parameters). Clusters were annotated based on the expression of top differentially expressed genes comparing each cluster to all remaining cells. Differential gene expression between groups was carried out using Mann–Whitney U tests and p-values were adjusted for multiple testing using the Bonferroni correction.

To identify progression-associated cell states, we performed a label-free differential abundance (DA) analysis using *DAseq* on the PCA embedding^82^. Pathway enrichment analysis of upregulated and downregulated DEGs was conducted using gprofiler2, querying the KEGG and Reactome (REAC) gene sets ^83^.

Trajectory analysis of the monocyte-to-macrophage transition was performed using *scVelo* and *CellRank2*^84,85^. RNA velocity was computed, and the resulting velocity pseudotime was incorporated into CellRank’s pseudotime kernel using a Generalized Perron Cluster Cluster Analysis estimator. To visualize gene expression dynamics along the transition towards LAM2 or LAM4, we fitted generalized additive models using CellRank-derived fate probabilities and visualized gene expression trends in a heatmap. Transcription factor activities were inferred using *decoupler*, training a univariate linear model on each transcription factor’s predicted regulon^86^. Pathway enrichment analysis of upregulated and downregulated DEGs

Cell–cell communication analysis was performed using *nichenetr* 2.2.0. For the *get_expressed_genes* function, the default expression threshold (pct = 0.1) was filtered to 0.05. DEG in receiver populations (MdC: comparing progressive to acute EAE) were filtered for pval-adj≤ 0.05 and log2fc ≥ 0.25, and the top 200 genes with the lowest adjusted p-values were retained. Ligand activity scores were computed using the Pearson correlation method instead of the default *aupr_corrected*, and the top 30 upstream ligands were selected. Ligands were assigned to cell types using a modified *assign_ligands_to_celltype* function, replacing the default mean + SD cutoff with a maximum-based approach (max(x) -0.00001) to exclude the “general” category. For visualization, ligand–target links were obtained using the *get_ligand_target_links_oi* function with the cutoff lowered from 0.04 to 0.0, ensuring that no low-weight links were discarded. These filtering yielded 162 predicted ligands.

### Serum GFAP and NfL Measurements

Serum NfL and GFAP from control (CFA-immunized) and EAE mice (acute and progressive phase) were measured in a blinded manner on the same day, using the Neurology 2-plex B assay (Quanterix, Billerica, MA) according to manufacturer’s instructions on the single molecule array SR-X Biomarker detection system. Samples were diluted by the factor 16 and measured in duplicates. Measurements that exceeded an intra-assay coefficient of variation (CV) of 20% were excluded. Two internal quality controls from the kit were included in each run with mean CV of 19.68% for low control and 13.03% for high control for sGFAP (concentrations: pg/ml) and 9.32 % for low control and 1.95 % for high control for sNFL (concentrations pg/ml).

### Immunohistochemistry (IHC)

CNS tissue samples were prepared for IHC as previously described in^12^. Spinal columns were dissected out and drop-fixed for 24 h in 4% paraformaldehyde (PFA) (Morphisto) at 4 °C followed by decalcification in RDFTM (Biosystems, 10-20% formic acid) for 48h followed by dehydration in graded alcohols. Transversal lumbar sections of the SC were dissected and embedded in paraffin. 3µm-thick tissue sections were cut on a microtome (Microm HM 360, Thermo Fisher). Serial sections were mounted on the same slide. Sections of the lumbar spinal cord segments were stained with a primary antibody directed to myelin basic protein (MBP; Merck Millipore1:800), Iba1 (Wako, 1:750) or a non-phosphorylated neurofilament SMI-32 (nNF; BioLegend, 1:4000). Secondary signal was detected using the horseradish peroxidase (HRP) method. Images were acquired using NanoZoomer 2.0-HT; Hamamatsu, Hamamatsu City, Japan or Zeiss Axio Scan. Z1 Slidescanner (Zen 2 software, blue edition) with 10x or 20x objectives. The scanned slides of each animal were quantitatively analyzed using the Visiopharm 2022.01.3.12053 software (Visiopharm, Hoersholm, Denmark).

To quantify demyelinated area and to count SMI-32*+* elements representing axonal damage, regions of interest (ROIs) corresponding to the white matter of the SC were manually drawn. Within these ROIs, the MBP positive area was calculated. Demyelinated area was defined as total white matter area of the SC circumference-MBP*+*%. We then report the average percentage demyelination of each mouse. The count of SMI-32*+* elements was performed and was calculated as positive elements per mm*2* of white matter. These analyses were performed in a blinded manner by expert pathology service at the Laboratory for Animal Model Pathology (LAMP) from the University of Zurich.

### Multiplex immunofluorescence

Sequential immunofluorescence analysis was performed using the COMET platform (Lunaphore Technologies). SC tissues were prepared for imaging similar to that described in^12^. Briefly, around 2mm of spinal cords were dissected and fixed for 24 h in 4% PFA and then decalcified in 0.5M EDTA solution (Axonlab) for five consecutive days both at 4°C. Spinal cords were then transferred to 30% sucrose in PBS for 48 h at 4°C, embedded in cryo-embedding medium (Medite) and frozen at −80°C for sectioning. The frozen lumbar SC were cryosectioned into 8-μm sections onto super frost glass slides and stored at −20 degrees. Slides were washed with PBS and simultaneously blocked and permeabilized using 10% NDS diluted in PBS-T (0.3% Triton (Sigma) diluted in PBS) for 1.5 hours at room temperature. Samples were loaded onto the COMET Stainer and processed according to the manufacturer’s instructions. Samples underwent iterative staining with primary antibodies, secondary antibodies, and subsequent imaging, followed by elution of the primary and secondary antibodies. All primary antibody incubations were performed for 8 minutes. A list of antibodies used can be found in (**Supplementary Table 4**). Lunaphore Viewer was used to view the images and remove background fluorescence. QuPath 0.6.0-rc3 was used to identify and annotate disrupted regions, and perform cell segmentation using Instanseg*86*, an embedding-based instance segmentation algorithm that uses DAPI and specified markers signals (CD11c, CD3, GFAP, Iba1, MBP, NeuN, P2RY12). After segmentation, each cell’s x and y coordinates and the median fluorescence signals from the nucleus and cell were exported for downstream R analysis.

### Multiplex immunofluorescence analysis

Before automated high-dimensional data analysis, multiplex imaging data were transformed using an inverse hyperbolic sine (arcsinh) function with a manually determined coefficient and normalized between 0 and 1. A preliminary round of unsupervised clustering using FlowSOM was conducted to clean the data and exclude unstained cells from downstream analysis. The resulting cells were annotated as homeostatic microglia (hMglia), Reactive astrocytes, oligodendrocytes, LAM cluster 4, DAM/MdCs 1, T cells, DAM/MdCs 2, neurons and LAM cluster 2 based on the markers (ARG-1, CD11c, CD3, Dectin-1, GFAP, GPNMB, Iba1, MBP, MerTK, NeuN, iNOS, P2RY12, TREM2).

To assess the spatial proximity of annotated cell clusters, we calculated k-nearest neighbor (kNN) based on neighborhood enrichment scores (Z-score). For each cell, spatial coordinates (x, y) and annotated cluster data were extracted. The FNN R package was used to identify the 10 nearest neighbors for each cell based on Euclidean distance. For every cluster pair, we counted how often a given source cluster was observed among the neighbors of a target cluster, generating a cluster-to-cluster neighborhood count matrix. Counts were normalized to the expected frequencies from the global cluster distribution, yielding an enrichment ratio (observed/expected). Values greater than 1 indicated that a cluster was found in the neighborhood of another cluster more often than expected by chance. Z-score or enrichment toward the LAM cluster 4 was extracted and further plotted.

### Analysis of publicly available datasets

Processed AnnData objects from Kukanja *et al*.^39^(Zenodo: 10.5281/zenodo.8037425) were downloaded and reanalyzed in Python 3.12 using scanpy 1.10.4 and squidpy 1.6.5. Mouse and human datasets were processed separately.

For the mouse dataset, samples were reclustered using the Leiden algorithm with a resolution of 3. To quantify LAM gene expression, we computed module scores using the following LAM marker set derived from differentially expressed genes in the NOD EAE scRNA-seq dataset and filtered for genes present in the spatial transcriptomics panel: *Anxa5, B2m, Cd24a, Cd44, Cd74, Cdkn1a, Crip1, Dusp1, Egr1, Egr2, Fcgr2b, Fos, Fosb, Gng11, H2-Aa, H2-Ab1, H2-Eb1, Hopx, Lgals1, Mrc1, Ms4a7, Mt1, Mt2, Pf4, S100a10, Sdc1, Sgk1*. Cluster 3, which displayed the highest median LAM module score, was annotated as LAMs. Neighborhood enrichment analysis was then performed using the squidpy neighborhood enrichment function on the “EAE_late” subset. This function computes enrichment by permutation testing and outputs both z-scores and enrichment counts. Lollipop plots were generated from neighborhood enrichment results for the LAM subset in the “EAE_late” stage.

For the human dataset, cells were subsetted based on original annotations to retain only immune populations: vascular_MP_1, vascular_MP_2, vascular_MP_3, vascular_T-cell, MP/MiGl_1, MP/MiGl_2, and T-cell. The immune subset was reclustered using the Leiden algorithm with a resolution of 2. LAM module scores were computed using the following gene set, filtered for presence in the Xenium panel: *ANXA1, APOE, APP, CAPG, CD14, CD68, CXCL14, CXCR4, FCER1G, GPNMB, HILPDA, IDH1, IL7R, ITGB2, MGST1, PSEN2, RFTN1, TGFBI, THBS1*. Cluster 15, with the highest median LAM module score, was annotated as LAM. This annotation was merged back into the full dataset (including non-immune cells), and neighborhood enrichment was computed on the “ms_active” subset. Lollipop plots were generated using neighborhood enrichment z-scores and counts for the LAM subset in the “ms_active” stage.

Processed snRNA-seq AnnData object from Lerma-Martin *et al.*^46^(GSE279183) was downloaded and converted to a Seurat object for further R analysis. Cells were subsetted based on original annotations to retain only myeloid populations: MG_CA, MG_Dis, MG_Homeo1, MG_Homeo2, MG_Homeo3, MG_NA, MG_Phago1 and MG_Rim. The myeloid subset was reclustered using the Leiden algorithm with a resolution of 2. The resulting 11 clusters were manually annotated as hMglia (0, 3), DAM ( 8, 4, 7, 10, 5, 6, 1), MdCs (2,11) and BAMs (9). BAMs were excluded from further analysis. Within the myeloid annotated clusters, LAM module scores were computed using the following gene set: *GPNMB, APOC4, APOE, CTSD, PSAP, LYZ2, CD63, TIMP2, ITM2B, SYNGR1, TREM2, LAMP1, FABP5, ACP5, HEXA, PLD3, CTSS, MS4A7, MERTK, CTSB, PLTP, SELENOP, TMEM37, GNGT2, ABCA1, CD68, FTL1, DHRS3, CDO1, LGALS3BP, LGMN, CD300A, CREG1, MAN2B1, C1QC, GDF15, CD300C2, GNS, AXL, GRN, LIPA, RGS1, APOBEC1, PLIN2, SNX5, CAMK1, F7, RGS10, SGK1, C1QA, CTSL, C1QB, H2-AA, PTMS, CEBPA, H2-EB1, H2-AB1, MAF, CD74, FABP4, APOC1, SAA3*.

### High dimensional analysis of FC data

Compensated and cleaned flow cytometry data were exported from FlowJo and analyzed in RStudio (version 4.4.0). High-dimensional analysis was performed as previously described^87^. Briefly, data were arcsinh-transformed using variable manually set co-factors and normalized using percentile normalization. Throughout the downstream analysis these arcsinh transformed and scaled expression values were utilized. UMAP was performed using the *umap* package to visualize cellular phenotypes in reduced dimensions in a random subset of the data. Clustering of leukocytes was conducted with the *FlowSOM* package, followed by metaclustering using the *ConsensusClusterPlus* package. Metaclusters were merged and annotated based on their median marker expression profiles. Heatmaps of marker expression were generated using the pheatmap package.

Principal component analysis of immune cell abundances and median marker expression per immune cell cluster was performed using the base *stats* package. Force-directed layouts for network visualizations representing intercluster relationships and trajectory structures were generated using the *grappolo* and *vite* packages. The layout algorithm for the graphs were automatically determined in *igraph* and visualized using *igraph* and *ggraph*.

To systematically retrieve immune features for which the progressive group demonstrated more extreme values than the acute group (both in relation to control mice), a chronification score 𝒄 was computed for the median expression of each marker in each cluster. This score was defined as the product of two Cohen’s *d* effect sizes: (i) between mice in the acute and progressive disease stages 𝛿*_AP_*_”_, and (ii) between control and progressive stages 𝛿*_NP_*:

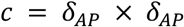

Cohen’s *d* effect sizes were calculated using the *effsize* package.

Finally, to visualize monocyte-to-macrophage differentiation trajectories, diffusion maps were computed using the *destiny* package and 3D visualizations were drawn using *rgl*. All other visualization were generated using *ggplot2*.

### Statistical Analysis

Data normality was assessed using Shapiro-Wilk test or Kolmogorov-Smirnov test. Based on normality results, group means were compared using either two-tailed, unpaired t-tests (for normally distributed data) or unpaired Mann-Whitney tests (for non-normally distributed data). Multiparametric analysis was performed using two-way analysis of variance (ANOVA), followed by Sidak’s multiple comparison tests. Corrections for multiple hypothesis testing were applied using the Benjamini-Hochberg method, and p-values are presented as adjusted p-values. Data are presented as mean ± standard error of the mean (SEM) or as mean ± standard deviation (SD), as reported in the respective figure legends. Data plotted in violin plots or box plots, depicts median and interquartile range. The variable n represents the number of biologically independent animals. P-values lower than 0.05 were considered statistically significant and are indicated by asterisks (*P < 0.05, **P < 0.01, ***P < 0.001). Numerical values and specific statistical methods used are reported in the respective figure legends or supplementary tables. Statistical analysis was carried out using R or GraphPad Prism 9 (GraphPad Software, Inc.).

### Declaration of generative AI and AI-assisted technologies in the writing process

During the preparation of this work the author(s) used ChatGPT in order to improve writing clarity. After using this tool/service, the author(s) reviewed and edited the content as needed and take(s) full responsibility for the content of the publication.

## Acknowledgements

We thank Yakine Raach, David Bamert and Can Ulutekin for technical support, and the Functional Genomics Center Zurich (FGCZ), in particular Hubert Rehrauer, and the Laboratory Animal Services Center (LASC) for providing equipment and services.

## Funding

This work has received funding from the Swiss National Science Foundation (Ambizione PZ00P3_193330 to S.M.), the Research Talent Development Fund zur Förderung des akademischen Nachwuchses (F-41302-40-01 to S.M.), Dr. Wilhelm Hurka Foundation (to S.M., D.D.F. and B.B.), the Swiss Multiple Sclerosis Society (Grant Number 2023-16 (to S.M.); Research Grant 2019/2020 (to S.M., D.D.F. and F.I.)), the Hartmann-Müller Foundation (Nr. 2758 to S.M.), the Novartis Foundation for Medical-Biological Research (Nr. 23C17 to S.M.) and the UZH Candoc grant (FK-22-040 to J.V-V.). F.I. was supported by the Koshland price and an EMBO postdoctoral fellowship (ALTF 723-2022). F.I. is recipient of an MSCA postdoctoral fellowship (101106452; STIC-GBM). F.I. received funding from the Deutsche Forschungsgemeinschaft (DFG) research grant number 441891347 (SFB1479-OncoEscape). J.S. received funding from the Nationaal MS Fonds (OZ2018-003).

## Author contributions

J.V-V., D.D.F and S.M. designed and performed all mouse experiments. F.I., J.V-V., S.M and P.C. analyzed and interpreted all data and prepared all figures. V.B., E.M., J.K. D.B, C.A., E.R., P.Z., and S.S., helped with experiments. A.L. performed the SIMOA experiments. C.C.H, H.J.W., J.H. and J.S. performed the post-mortem human meningeal studies, H.V, M.M. and P.J. helped with histology analysis. S.M., F.I., B.B., D.D.F., M.G, S.T., M.H. and B.S. provided scientific and intellectual input. F.I., J.V-V. and S.M. wrote and P.Z., D.D.F., S.T., M.H., M.M., H.J.W., J.H., J.S. and B.S. edited the manuscript. S.M. and F.I., supervised and acquired funding for the study. All authors approved the paper.

## Data availability

Raw spectral flow cytometry data and scRNA seq data will be made publicly available in a repository upon publication.

## Code availability

Code for the analysis of cytometry and scRNA seq data will be uploaded into a GitHub repository upon publication.

## Competing interest declaration

The authors declare no competing interests.

---

**Figure S1:**
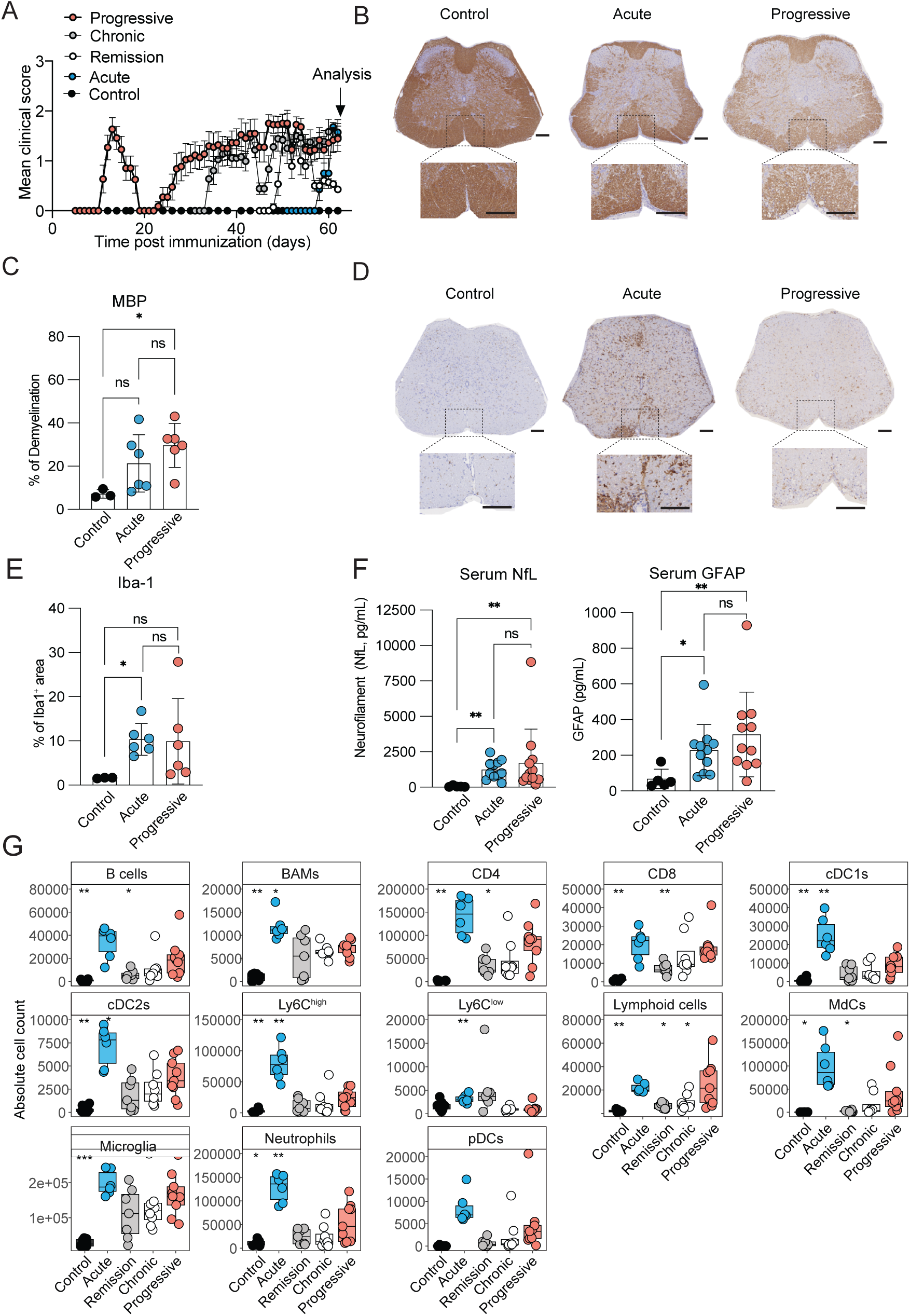
Progressive neuroinflammation is characterized by increased tissue damage and decreased immune cell infiltration. (**A**) Schematic of staggered immunization to analyse immune cell changes across progressive neuroinflammation. (**B**) Representative myelin base protein (MBP) staining and (**C**) the average percentage of demyelination in white matter in lumbar spinal cord (SC) sections from the depicted disease stages. (One-way ANOVA and Tukey’s multiple comparison; control vs progressive *P = 0.0316). (**D**) Representative Iba1 staining and (**E**) the average percentage of Iba1^+^ are in lumbar SC sections from the depicted disease stages. (One-way ANOVA and Tukey’s multiple comparison; control vs acute *P = 0.0294). For (**B** and **D**) Scale bar of 100uM. (**C-E**) Data from control (unimmunized, n=3, m/f), and mice with acute (n=6, m/f) and progressive (n=6, m/f) EAE. Data pooled from two independent experiments. (**F**) The average neurofilament (NfL) (left) and glial fibrillary acidic protein (GFAP) (right) serum levels. (Kruskall-Wallis ANOVA and Dunn’s multiple comparison; control vs acute **P=0.0029, control vs progressive **P = 0.0065 (left) and Dunn’s multiple comparison; control vs acute *P=0.0428, control vs progressive **P = 0.0099 (right)). Data from control (n=5, m/f), acute (n=11, m/f) and progressive (n=12, m/f) mice. Data pooled from three independent experiments. (**G**) Absolute cell counts from CNS leukocytes detected in NOD CNS across the different disease stages. Unpaired t-test; acute vs progressive: CD4 *P = 0.029, neutrophils **P = 0.001, Ly6C^hi^ **P = 0.002, Ly6C^low^ **P = 0.001, cDC1s **P = 0.008, cDC2s *P = 0.013, BAMs **P = 0.008; Chronic vs progressive: lymphoid cells *P = 0.044; Control vs progressive: lymphoid cells **P = 0.007, CD4 ***P < 0.001, CD8 ***P < 0.001, B cells **P = 0.007, neutrophils *P = 0.012, MdCs *P = 0.041, Ly6C^hi^ **P = 0.005, cDC1s **P = 0.003, cDC2s **P = 0.001, BAMs ***P < 0.001, microglia ***P < 0.001 and remission vs progressive: lymphoid cells *P = 0.019, CD4 *P = 0.015, CD8 **P = 0.003, B cells *P = 0.029, MdCs *P = 0.049. Data from control (CFA-immunized, n=8, m/f), acute (n=6, m/f), remission (n=7, m/f), chronic (n=8, m/f) and progressive (n=9, m/f). Data are representative for one out of two independent experiments.

**Figure S2:**
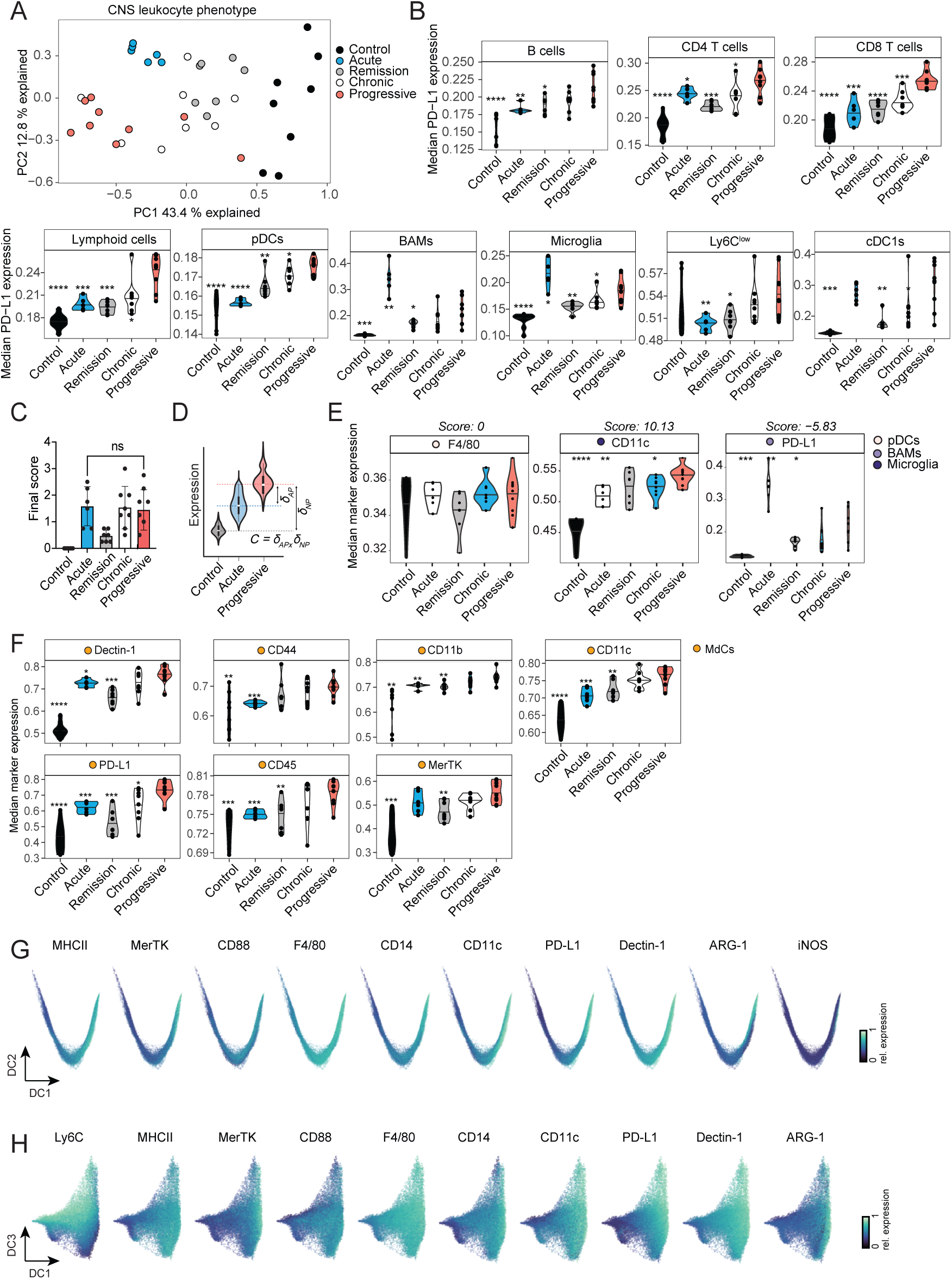
Full-spectrum cytometry analysis reveals altered monocyte-to-macrophage transition during progressive neuroinflammation. (**A**) PCA graph of CNS leukocyte phenotype in NOD EAE disease stages. Dots represent individual mice, and color represents the disease stage. (**B**) Violin plots showing the median PD-L1 expression in the different leukocyte clusters across NOD disease stages. Unpaired t-test; progressive vs chronic: CD4 *P = 0.035, CD8 ***P < 0.001, cDC1s *P = 0.046, lymphoid cells *P = 0.013, , microglia *P = 0.044, pDCs *P = 0.045; progressive vs remission: B cells *P = 0.014, BAMs *P = 0.011, CD4 ***P < 0.001, CD8 ****P < 0.0001, cDC1s **P = 0.001, cDC2s ***P < 0.001, Ly6C^hi^ **P = 0.002, Ly6C^low^ *P = 0.013, lymphoid cells ***P < 0.001, , microglia **P = 0.003, neutrophils **P = 0.006, pDCs **P = 0.004; Progressive vs acute: B cells **P = 0.003, BAMs **P = 0.002, CD4 *P = 0.030, CD8 ***P < 0.001, Ly6C^low^ **P = 0.005, lymphoid cells ***P < 0.001, microglia *P = 0.042, pDCs ****P < 0.0001; progressive vs control: B cells ****P < 0.0001, BAMs ***P < 0.001, CD4 ****P < 0.0001, CD8 ****P < 0.0001, cDC1s ***P < 0.001, cDC2s ***P < 0.001, Ly6C^hi^ **P = 0.001, lymphoid cells ****P < 0.0001, MdCs ****P < 0.0001, microglia ****P < 0.0001, neutrophils **P = 0.004, pDCs ****P < 0.0001.(**C**) Final score from mice analysed. (**D**) Chronification scores were computed by multiplying the Cohens D effect size of median marker expression between the progressive and acute stage with the Cohens D effect size of median marker expression between the progressive stage and control mice. (**E**) Example of chronification scores: F4/80 in pDCs, score:0 (left); CD11c in microglia, score:10.13 (middle); and PD-L1 in BAMS, score:-5.83 (right). (**F**) Violin plots of median marker expression in MdCs. Unpaired t-test; progressive vs chronic: *P = 0.047 (PD-L1); progressive vs remission ***P < 0.001 (Dectin-1), **P = 0.009 (CD45), **P = 0.006 (CD11c), ***P < 0.001 (PD-L1), **P = 0.004 (MerTK); progressive vs acute *P = 0.027 (Dectin-1), ***P < 0.001 (CD45), ***P < 0.001 (CD44), **P = 0.003 (CD11b), ***P < 0.001 (CD11c), ***P < 0.001 (PD-L1); control vs progressive ****P < 0.0001 (Dectin-1), ***P < 0.001 (CD45), **P = 0.008 (CD44), **P = 0.004 (CD11b), ****P < 0.0001 (CD11c), ****P < 0.0001 (PD-L1), ***P < 0.001 (MerTK)). (**G-H**) Diffusion maps of marker relative expression in (**G**) DC2 vs DC1 and (**H**) DC3 vs DC1 in MdCs. (**A-H**) Data from control (CFA-immunized, n=8, m/f), and mice with acute (n=6, m/f), remission (n=7, m/f), chronic (n=8, m/f) and progressive (n=9, m/f) EAE. Data are representative for one out of two independent experiments.

**Figure S3:**
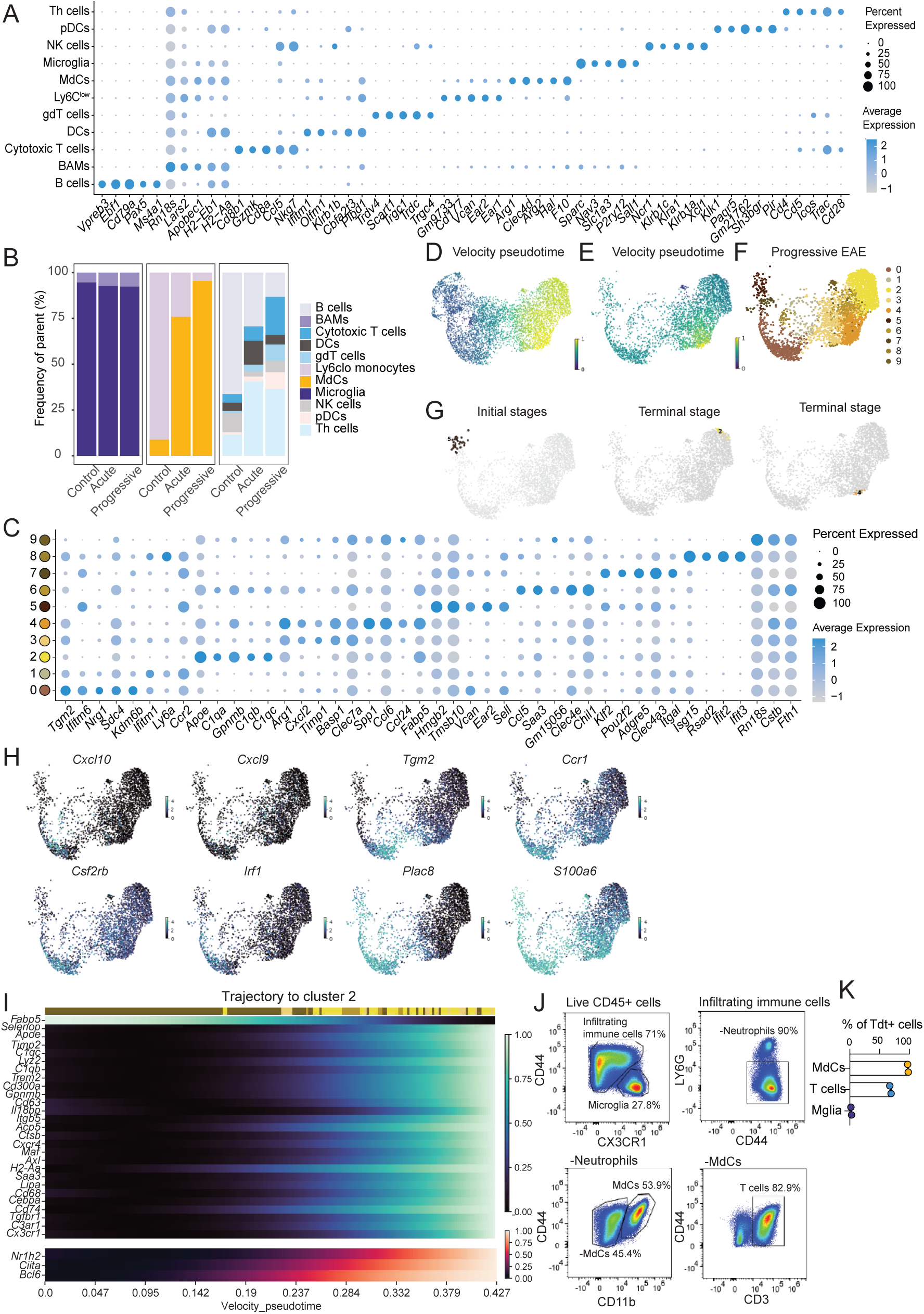
Single-cell RNA sequencing confirms chronic monocyte-to-phagocyte transition as a hallmark of progressive disease. (**A**) Dot plot of marker average expression in the detected immune cell clusters in NOD CNS by scRNAseq. (**B**) Bar plot of frequency of parent of resident phagocytes, infiltrating phagocytes and other leukocytes across control, acute and progressive disease stages. (**C**) Dot plot of marker average expression in the subset MdCs clusters.(**D-E**) UMAP overlay of velocity pseudotime on subset MdCs from (**D**) merged control, acute and progressive or (**E**) only progressive EAE. (**F**) UMAP of MdCs subclusters in progressive disease stage. (**G**) UMAP overlay of predicted initial stages (left), and terminal stages (middle and right) predicted using Cellrank. (**H**) UMAP overlay of *Cxcl10, Cxcl9, Tgm2, Ccr1, Csf2rb, Irf1, Plac8* and *S100a6* in MdCs in progressive EAE. (**I**) Dot plot of marker average expression in MdCs cluster 0, 2 and 4 from progressive stage. (**J**) Heatmap displaying smoothed gene expression dynamics (top) or transcription factor regulon activities (bottom) that correlate with absorption probability for MdCs subcluster 2. Transcription factor activity has been estimated based on target transcript expression using Decoupler. (**K**) Representative pseudocolor dot plots depicting gating strategy of- and (**L**) percentage of TdT^+^ cells found in microglia, MdCs and T cells analysed from acute (d=14, n=2) *Ccr2^C^*^reERT2/+^ *Rosa26*^Ai14/+^ x NOD mice CNS. For (**A-J**) data shown corresponds to control, acute, and progressive stages (n=1, f). For (**A,C,I**) Circle size is depicting % of normalized expression detected per disease stage and color is depicting expression level as average expression.

**Figure S4:**
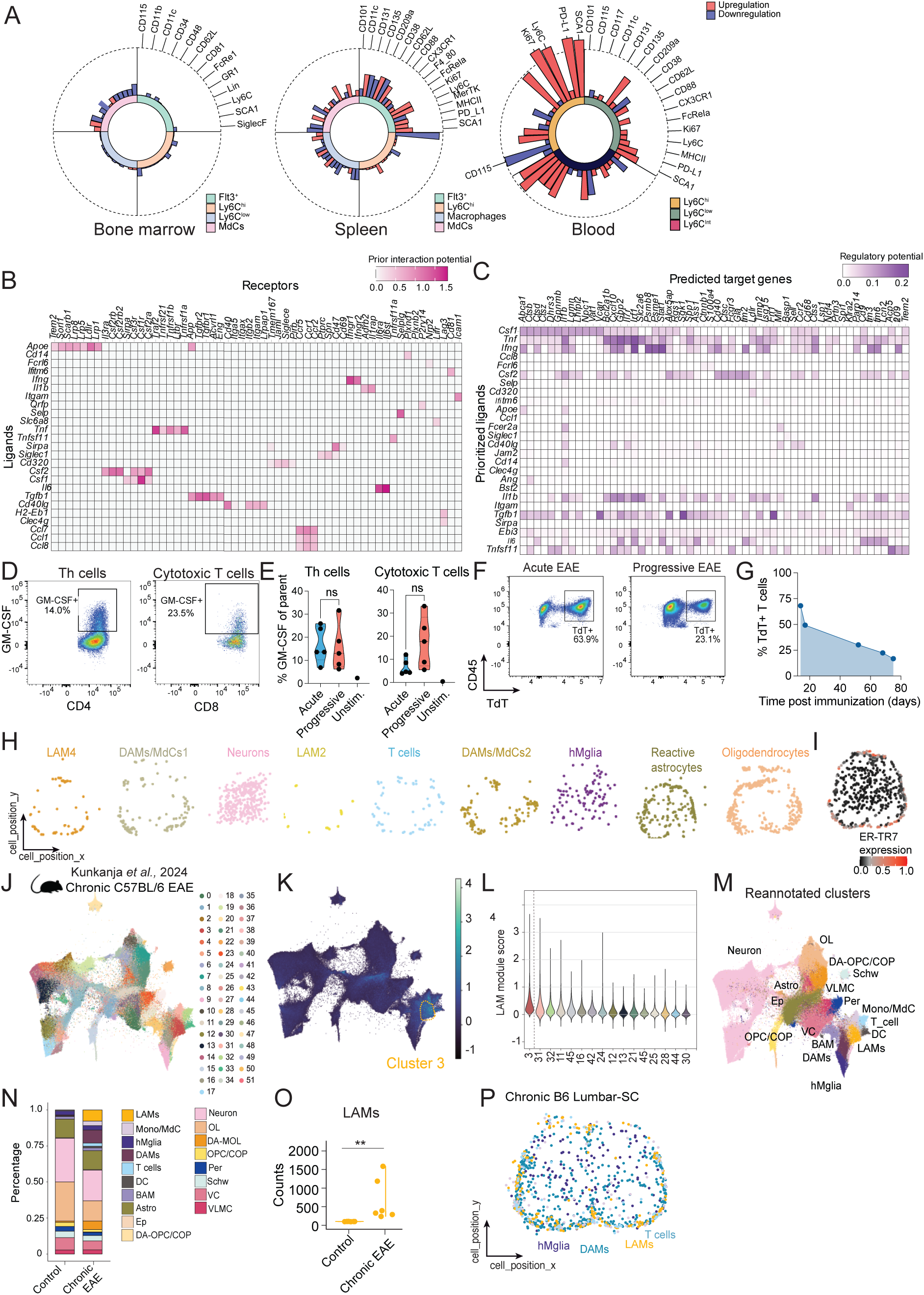
Progression-associated MdCs are imprinted locally in the leptomeninges. (**A**) Radar plots showing changes in the marker expression of monocyte subsets found in bone-marrow (BM), spleen and blood from progressive compared to acute stage. Values correspond to −log10 of adjusted p value. Inner circle color annotations denote each respective subset. Bar color denotes the upregulation or downregulation of the respective marker. Marker order is consistent across subsets. Dashed line denotes −log10(0.05) cutoff for p value. Heatmap of (**B**) prior interaction potential and (**C**) predicted target genes from the Nichenet analysis. (**D**) Representative pseudocolor FC-dotplots depicting gating strategy of- and (**E**) percentage of GM-CSF^+^ cells found in PMA/Ionomycin restimulated T cells isolated from the CNS of mice with acute (n=5, m/f) and progressive (n=5, m/f) EAE. Last dot in (**E**) indicates technical non-stimulated control (n=1). (**F**) Pseudocolor FC-dotplots depicting positive Ai14 (TdT^+^) T cells in acute and progressive stages. (**G**) Percentage of TdT^+^ labelling in T cells over the course of progressive neuroinflammation. For (**F-G**) data corresponds to: day 14 n=2 m/f; day 17 n=1 f; day 52 n=2 f; day 68 n=1 m; and day 75 n=3 m/f. (**H**) Cell position (x and y) of cell clusters identified using Lunaphore COMET^TM^ multiplex microscopy platform in lumbar progressive SC sections. (progressive, n=4 m/f). (**I**) Representative overlay of ER-TR7 expression in progressive SC section. (**J-P**) Reanalysis of chronic EAE data from Kukanja *et al*. (**J**) UMAP of clusters detected. (**K**) UMAP overlay of the gene signature (cluster 3 is highlighted) and (**L**) violin plot of the module score overlay of LAMs. (**M**) UMAP and (**N**) bar plot of annotated cell clusters. (**O**) Violin plot depicting LAM counts in control and chronic C57BL/6 EAE (Mann-Whiteny test, **P= 0.0022) reanalyzed from Kukanja *et al.*. (**P**) Representative overlay of T cells, LAMs and DAMs in SC lumbar section.

**Figure S5:**
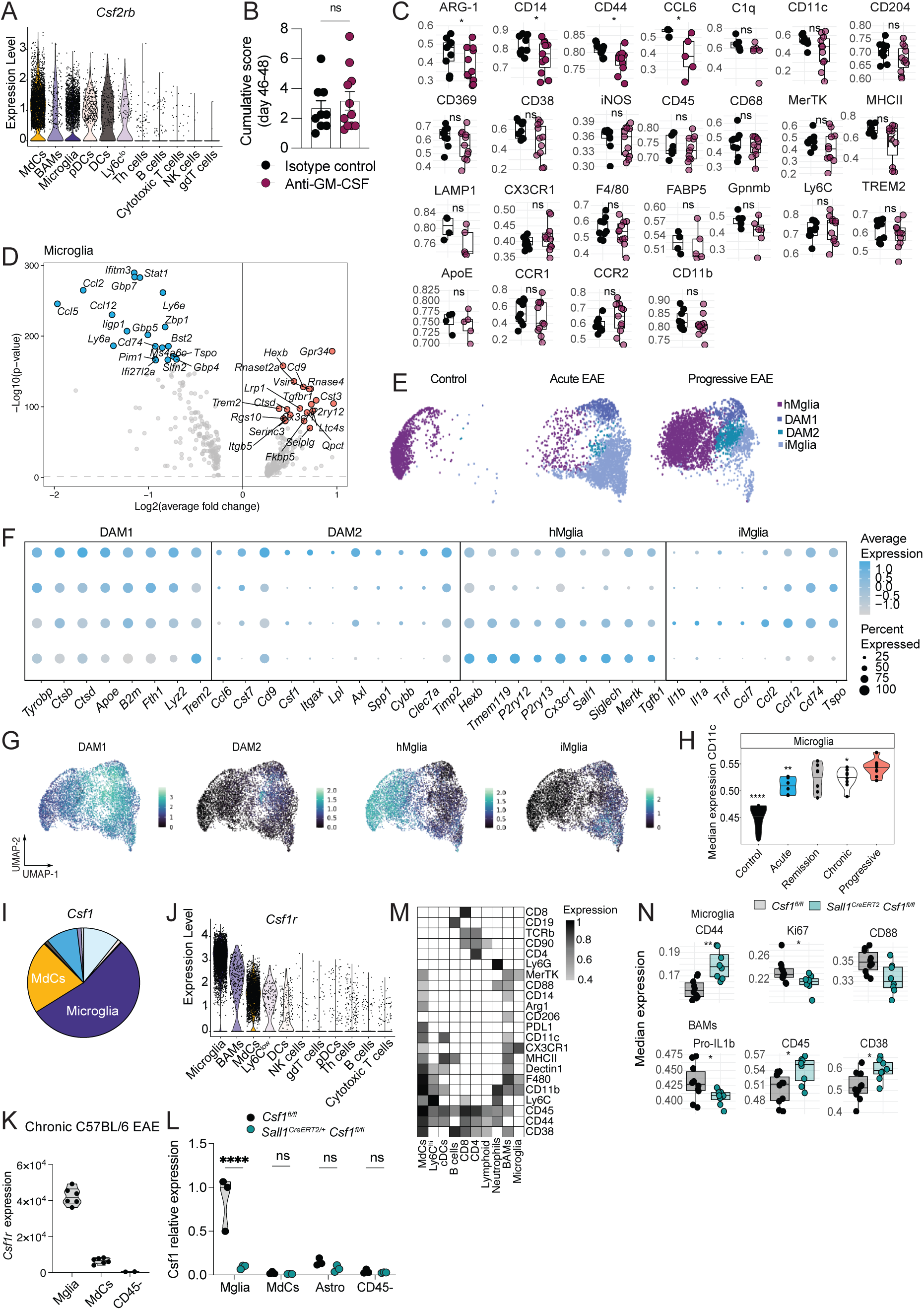
DAM2-like microglia are present in both models of progressive and chronic MS neuroinflammation. (**A**) Violin plot of *Csf1r* expression across the detected CNS immune cell clusters in progressive NOD EAE. (**B**) Cumulative clinical score of short-term treated isotype control treated (n=9, m/f) and anti-GM-CSF treated (n=11, m/f) mice. Unpaired two-tailed t-test. (**C**) Boxplot of marker media expression in MdCs from short term treated isotype control treated (n=9, m/f) and anti-GM-CSF treated (n=11, m/f) mice. Pooled data from two independent experiments analyzed using a linear regression model: *P = 0.0202 (ARG-1); *P < 0.026 (CD44); *P = 0.0487 (CD14) and unpaired t-test *P=0.048 (CCL6). (**D**) Differentially expressed genes (DEGs) in microglia when comparing progressive to acute stage of the disease. Blue dots indicate markers upregulated in acute and red dots markers upregulated in progressive. (**E**) UMAP of microglia subclusters detected by scRNAseq and split by disease stages. (**F**) Dotplot of marker average expression in the subset microglia clusters grouped by DAM1, DAM2, hMglia (homeostatic microglia) and iMglia (inflammatory microglia) markers. (**G**) UMAP overlay of the gene signatures DAM1, DAM2, hMglia and iMglia. (**H**) Violin plot of the median CD11c expression (flow cytometry) in microglia across NOD disease stages. Unpaired two-tailed t-test: control vs progressive ****P<0.0001; acute vs progressive **P=0.0016; chronic vs progressive *P=0,02999. Data from control (unimmunized, n=8, m/f) and mice with acute (n=6, m/f) remission (n=7, m/f), chronic (n=8, m/f) and progressive (n=9, m/f) EAE. Data from one representative cohort (**I**) Pie chart depicting the distribution of *Csf1* expressing and (**J**) violin plot of *Csf1r* expression across leukocyte clusters in progressive NOD CNS scRNAseq data. (**K**) Median of *Csf1* expression in FACS-sorted microglia, MdCs and CD45 negative cells from chronic C57BL/6 EAE (d24), (n=6 for Mglia and MdCs, n=2 for CD45^-^ fraction, m/f). (**L**) *Csf1* relative expression in Mglia, MdCs, astrocytes (Astro) or CD45--remaining fractions FACS-sorted from the inflamed CNS of *Sall1^+/+^ Csf1*^fl/fl^ (n=3, f) and *Sall1^CreERT2/+^ Csf1*^fl/fl^ (n=3, f) mice. (**M**) Heatmap showing the median marker expression of the identified immune cell clusters. (**N**) Median fluorescence intensity (MFI) analysis of activation markers in microglia and BAMs from the inflamed CNS of *Sall1^+/+^ Csf1*^fl/fl^ (n=9, m/f) and *Sall1^CreERT2/+^ Csf1*^fl/fl^ (n=8, m/f) mice. Unpaired t-test: microglia (**P=0.0015 (CD44), *P=0.293 (Ki67), *P=0.028 (CD88)) and BAMs (*P=0.0324 (pro-IL1b), *P=0.015 (CD45), *P=0.0115 (CD38)).

**Figure S6:**
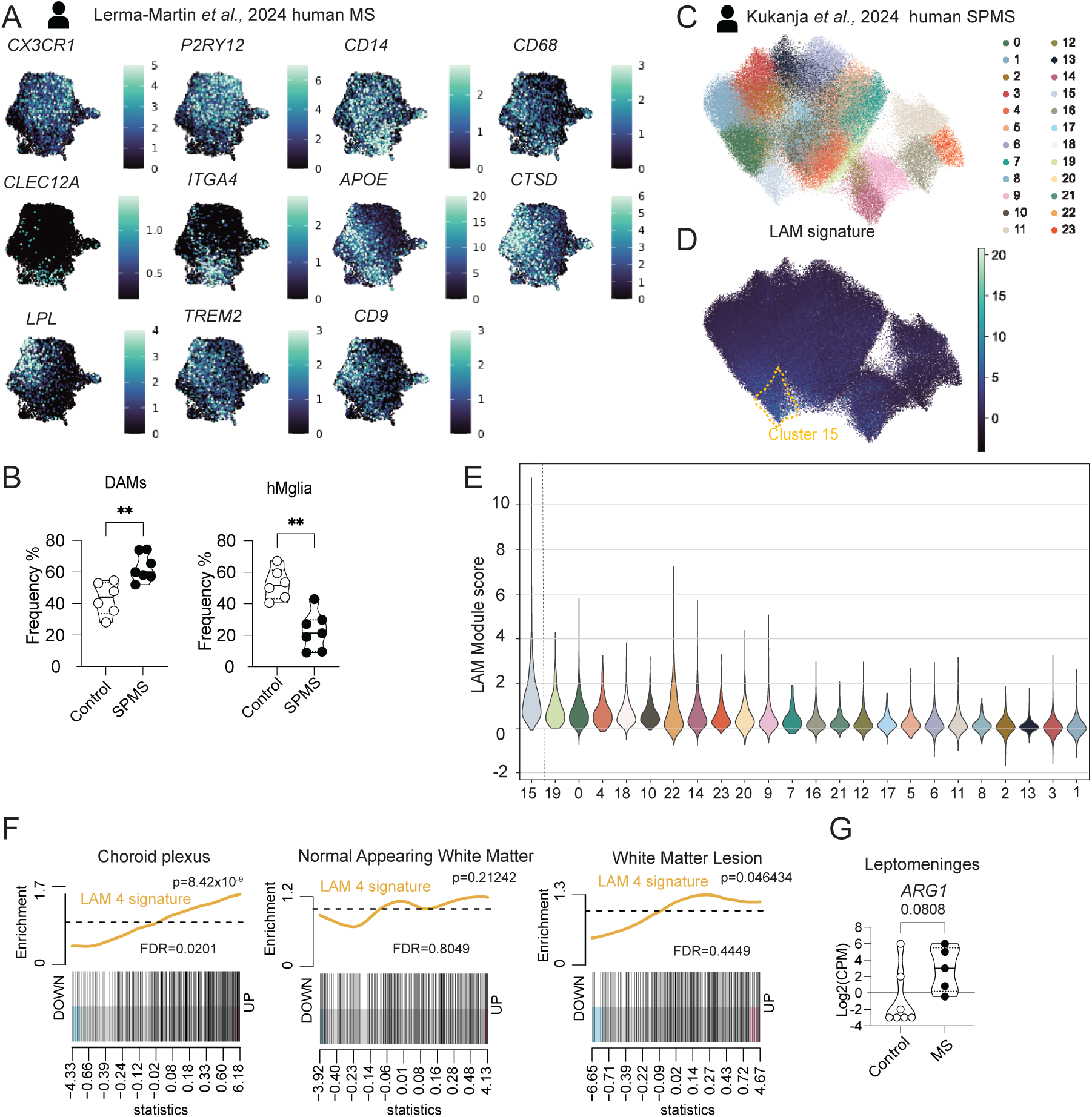
LAMs and their local cues are present in SPMS patients. (**A**) UMAP overlay of the genes used to cluster the phagocyte subsets detected human subcortical tissue of control and MS patients; reanalyzed from Lerma-Martin *et al.*^46^. (**B**) Frequency of disease-associated microglia (DAMs) and homeostatic microglia (hMglia) found in control and MS patients. (Mann-Whitney test, **P= 0.0047 for DAMs , **P=0.0023 for hMglia, data from control (n=6) and MS patients (n=7)) (**C-E**) Reanalysis of human SPMS data from Kukanja *et al.*^39^ (**C**) UMAP of myeloid subclusters (**D**) UMAP overlay of the LAM gene signature (cluster 15 highlighted) and (**E**) violin plot of the LAM module score per subcluster. (**F**) GSEA plot based on the LAM cluster 4 signature enriched analysis in myeloid cells from MS vs control patients: choroid plexus (left), normal appearing white matter (middle), and white matter lesion (right). (**G**) Log2 counts per million of Arginase-1 (ARG-1) detected in control and MS patient meninges (Mann-Whitney test P= 0.0808). Data reported on **Supplementary Table 3**. (**F**-**G**) Data from control (n=7) and MS patients (n=5).

**Supplementary Table 1. DEGs in MdCs Progressive vs Acute.**

**Supplementary Table 2. DEGs in Microglia Progressive vs Acute.**

**Supplementary Table 3. Antibodies used in full-spectrum flow cytometry and cell sorting.**

**Supplementary Table 4. Antibodies used in multiplex imaging.**

**##Supplementary Table 5. RNA enrichment analysis of postmortem brain samples.**

## Notes

### Competing Interest Statement

The authors have declared no competing interest.

